# Thyroid hormone rewires cortical circuits to coordinate body-wide metabolism and exploratory drive

**DOI:** 10.1101/2023.08.10.552874

**Authors:** Daniel R. Hochbaum, Alexandra C. Dubinsky, Hannah C. Farnsworth, Lauren Hulshof, Giona Kleinberg, Amanda Urke, Wengang Wang, Richard Hakim, Keira Robertson, Canaria Park, Alyssa Solberg, Yechan Yang, Caroline Baynard, Naeem M. Nadaf, Celia C. Beron, Allison E. Girasole, Lynne Chantranupong, Marissa Cortopassi, Shannon Prouty, Ludwig Geistlinger, Alexander Banks, Thomas Scanlan, Michael E. Greenberg, Gabriella L. Boulting, Evan Z. Macosko, Bernardo L. Sabatini

**Affiliations:** Howard Hughes Medical Institute, Department of Neurobiology, Harvard Medical School; Society of Fellows, Harvard University; Broad Institute of MIT and Harvard; Department of Biomedical Informatics, Harvard Medical School; Department of Medicine, Beth Israel Deaconess Medical Center, Harvard Medical School; Department of Neurobiology, University of Massachusetts Chan Medical School; Center for Computational Biomedicine, Harvard Medical School; Department of Chemical Physiology & Biochemistry, Oregon Health & Science University School of Medicine; Department of Neurobiology, Harvard Medical School; Department of Psychiatry, Massachusetts General Hospital

**Author notes:** These authors contributed equally, are ordered alphabetically, and can list themselves as 2^nd^ author on their C.V.s.

## Abstract

Animals adapt to varying environmental conditions by modifying the function of their internal organs, including the brain. To be adaptive, alterations in behavior must be coordinated with the functional state of organs throughout the body. Here we find that thyroid hormone— a prominent regulator of metabolism in many peripheral organs— activates cell-type specific transcriptional programs in anterior regions of cortex of adult mice via direct activation of thyroid hormone receptors. These programs are enriched for axon-guidance genes in glutamatergic projection neurons, synaptic regulators across both astrocytes and neurons, and pro-myelination factors in oligodendrocytes, suggesting widespread remodeling of cortical circuits. Indeed, whole-cell electrophysiology recordings revealed that thyroid hormone induces local transcriptional programs that rewire cortical neural circuits via pre-synaptic mechanisms, resulting in increased excitatory drive with a concomitant sensitization of recruited inhibition. We find that thyroid hormone bidirectionally regulates innate exploratory behaviors and that the transcriptionally mediated circuit changes in anterior cortex causally promote exploratory decision-making. Thus, thyroid hormone acts directly on adult cerebral cortex to coordinate exploratory behaviors with whole-body metabolic state.

## Introduction

In response to varying environmental conditions, such as changes in food availability and seasonal variation in weather, animals coordinate alterations in the function of their organs with changes in their patterns of behavior. For example, snakes that undergo long periods of fasting punctuated by large meals will, after swallowing their prey, reassemble digestive organs and seek atypical habitats in which they become largely immobile^1^. Similarly, animals that seasonally hibernate drastically change organ function, metabolism, and body temperature, while also engaging a dormant behavioral state, in which they suppress both thirst and feeding drives^2^. The coordination of peripheral (i.e., outside of the brain) organ function with changes in behavior is therefore often adaptive, enabling animals to survive in fluctuating and uncertain environments.

Signals to coordinate these processes likely include circulating hormones that are produced in and act on peripheral organs but also enter the brain where they act on receptors expressed by diverse classes of brain cells, including neurons, glia, immune, and vasculature-associated cells. Hormones are often components of essential homeostatic (i.e., negative feedback) control systems mediated by the hypothalamus— for example, leptin produced by adipocytes acts via hypothalamic circuits to reduce food consumption and increase energy expenditure, thus stabilizing body fat composition^3,4^. However, circulating hormones have additional effects in the brain beyond homeostatic control that modulate behavior and cognition. For instance, leptin influences visual cortices to control the degree to which brain cells consume energy to precisely code visual information, triggering expenditure of cellular energy supplies, when abundant, to improve visual perception^5^. Similarly, female sex hormones essential for ovarian and menstrual cycles as well as for fetal development enter the brain and induce nest building and other behaviors that need to be coordinated with pregnancy^6,7^. Thus, hormones that are generated and act peripherally can have non-homeostatic effects on the brain to coordinate changes in behavior with the remodeling of peripheral organs.

One hormone that affects both the periphery and the brain is thyroid hormone, which is produced in the thyroid gland and acts on many tissues to regulate metabolism and function. For instance, thyroid hormone stimulates lipolysis and fatty acid metabolism in the liver^8^, kindles thermogenesis in adipose tissue^9^, and increases energy expenditure, fast contractility of muscle fibers, and glycolysis in skeletal muscles^10,11^. Thyroid hormone also enters the brain and acts via the hypothalamic-pituitary-thyroid (HPT) axis to form a classic homeostatic feedback loop by which thyroid hormone inhibits its own production and stabilizes its circulating levels^12,13^. Thyroid hormone also acts on hypothalamic neurons to regulate body temperature, food intake, and weight^14,15^ (among other functions^16,17^). These observations in animals are consistent with changes in body temperature and weight in humans with pathologically low or high thyroid levels characteristic of hypo- or hyper-thyroidism respectively^18,19^.

Seasonal fluctuations in thyroid hormone levels are pronounced in many mammalian species^20^. For instance, thyroid hormone levels surge in grey mouse lemurs from Madagascar during their resource abundant wet season. As a result, these small primates increase their metabolic rate by increasing their caloric intake four-fold and consuming twice as much oxygen^21^. In addition, these animals undergo dramatic changes in behavior. They expand their home territory in synchrony with increasing thyroid hormone levels, spending more time during the wet season awake, foraging for food, and searching for mates^22,23^. However, this shift incurs additional risk as these exploratory behaviors leave animals more prone to predation, and as a result their mortality rate increases during this season^24^. Similar correlations between thyroid levels and exploratory-like behaviors have been observed in many species in the wild^17,25–28^.

Further evidence that thyroid hormone drives changes in exploratory behaviors comes from humans with dysregulated thyroid hormone levels. In addition to extreme changes in metabolism, individuals with hypo- and hyper-thyroidism often exhibit psychiatric symptoms ranging from depression in hypothyroidism to mania in hyperthyroidism^29–32^. These symptoms can be thought of as the pathological extremes of an axis of normal variation in exploratory behavior, with mania characterized as an over-expression of exploration relative to the risks present in the environment^33,34^, and depression characterized as a neglect of exploratory behaviors despite the presence of rewards, and absence of risks^35,36^. Collectively, evidence from animals and humans raise the possibility that thyroid hormone, in addition to changing metabolic state, could also drive the expression of exploratory behaviors.

The effects of thyroid hormone are mediated by thyroid hormone receptors (THRs), which are ligand-gated transcription factors that bind the active form of thyroid hormone, triiodothyronine (T3), with high affinity^12^. Unlike many steroid hormone receptors that translocate to the nucleus upon binding their ligand, unliganded THRs are bound to DNA and recruit co-repressors. Binding of T3 causes remodeling of chromatin resulting in the dissociation of co-repressors and recruitment of transcriptional activators^37^. Thus, THRs participate in active repression of target gene transcription in the absence of T3, and active transcription in the presence of T3. THRs are abundant not only in the adult hypothalamus, but also across cerebral cortex in both rodents^38,39^ and humans^40^. Although thyroid signaling is critical for proper cortical development^41,42^, its function in the adult cortex is unknown. Anterior cortical structures such as secondary motor cortex (M2), which are involved in higher order control of both goal-oriented and exploratory actions and decision-making, express THRs, and could therefore mediate potential thyroid-dependent changes in exploratory behaviors. Here we find that thyroid hormone directly remodels anterior cortical circuits through engagement of local thyroid-dependent transcriptional programs to coordinate exploratory-like behaviors with changes in body-wide metabolic state.

## Results

### Characterization of T3-modulated metabolism and behavior

Our goal was to identify thyroid-hormone dependent changes to neural circuits that underlie the alterations in exploratory behaviors observed in animals, including humans and many other species in the wild, with changes in levels of T3. We hypothesized that T3 directly affects cortical brain areas, such as M2, which express THR receptors and are known to be involved in higher order control of both goal-oriented and exploratory behaviors. Therefore, we first developed a T3-delivery paradigm that induced transcriptional changes in M2 of C57BL6/J mice and examined the utility of these mice for detecting T3-induced behavioral changes.

T3 was administered to adult mice via twice-daily intraperitoneal (IP) injections and comparisons were made to animals treated with vehicle alone (**Fig. 1a**, **Methods**). We used *Ier5* expression, a thyroid-responsive gene in developing neocortex^43^, as a readout of T3 entrance into, and action on, the brain. Expression of *Ier5* in frontal cortex increased with the concentration of T3 administered, saturated at doses of ∼0.1 ug/g (**Supp. Fig. 1a**), and was observed within an hour of treatment (**Supp. Fig. 1b**). T3 induced expression of several known thyroid-responsive genes (TRGs) throughout the layers of cortex (**Fig. 1b, Supp. Fig. 1c**), consistent with the presence of functional thyroid receptors in cortical tissue^38,39^. Sobetirome, a thyroid hormone mimetic with poor permeability across the blood brain barrier^44,45^, did not induce TRGs in cortex but did induce canonical TRGs in the liver (**Supp. Fig. 1d**). These observations are consistent with the induction of transcriptional programs in adult cortex by brain-penetrant thyroid hormone.

**Figure 1.**
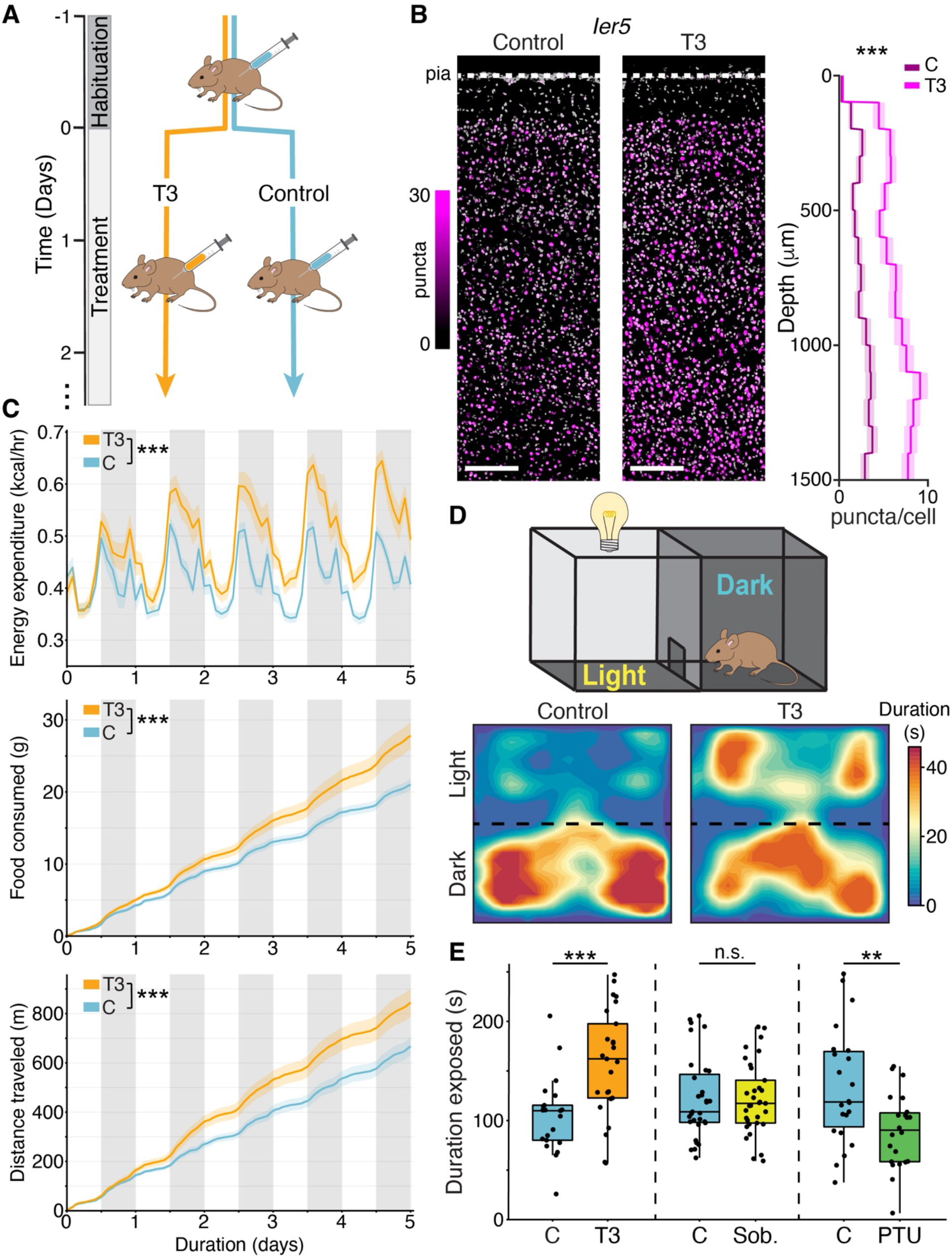
Brain-penetrant T3 induces transcription in cortex and modulates spontaneous exploratory behaviors. A) Schematic of experimental paradigm. Mice were habituated to IP injections with vehicle alone for 2 days prior to treatment. Mice were then divided into T3 and Control cohorts, with twice daily injections of T3 or vehicle, respectively. All transcriptional assays were conducted after 2.5 days of treatment, and behavioral assays at 3.5 days of treatment, unless otherwise noted. B) Fluorescence in situ hybridization (FISH) of the TRG *Ier5* in secondary motor cortex (left: control condition, middle: T3 condition). Scale bars = 200 μm. Right: summary of *Ier5* expression as a function of cortical depth. *Ier5* is upregulated by T3 across all layers of cortex (***p = 0, Wilcoxon rank-sum test comparing treatment effect across entire cortical depth; Control: n = 5494 cells; T3: n = 6061; 2 mice per condition). Central line indicates the mean, and shaded area are 95% confidence intervals of the mean. C) Home-cage indirect calorimetry performed over 5 days of treatment. Using generalized linear mixed models, we found that energy expenditure, food consumption, and distance traveled all significantly increased with T3 treatment over time (p < 10^-4^, likelihood ratio test, n = 16 control, n = 15 T3-treated mice). Central line indicates the mean, and shaded area the standard error of the mean. D) Top: Schematic of the light-dark preference assay (LD). Bottom: spatial heatmaps of the duration an example control (left) and T3-treated (right) mice occupied each area of the LD box. E) The duration mice stayed in the light-exposed zone of the LD assay increased with T3 (left) (p < 10^-3^, Wilcoxon-rank sum, n = 21 control, n = 25 mice), was unaffected by sobetirome (middle) (p = 0.87, Wilcoxon-rank sum, n = 31 control, n = 33 mice), and decreased with PTU (right) (p = 0.007, Wilcoxon-rank sum, n = 22 control, n = 22 mice). In these box plots the central line indicates the median and box and whiskers represent quartiles 1-4. For all panels: n.s. = not significant, ** = p < 0.01, *** = p < 0.001.

We used a home-cage indirect calorimetry system to survey the physiological processes affected by T3. T3 increased energy expenditure, food intake, and locomotion over the experimental time course (**Fig. 1c**; p < 10^-3^, likelihood ratio test). T3-treated animals also had elevated body temperatures (**Supp. Fig. 1e**), consistent with a heightened metabolic rate. Notably, the observed changes in physiology and coarse behaviors mirror effects of hyperthyroidism reported in humans, including hyperactivity.

To examine if increased thyroid hormone levels promote exploratory-like behaviors, we used a light-dark preference assay in which mice were placed in a box that contained both sheltered (dark) and light-exposed areas (**Fig. 1d)**. Mice naturally find the light-exposed region aversive and rarely explore this zone, instead spending most of their time in the sheltered, dark region. Mice treated with T3 (3.5 days) spent more time exposed in the light than vehicle-treated animals (**Fig. 1d-e**). The magnitude of the effect depended on T3 concentration but remained significantly elevated even at the lowest T3 levels tested (**Supp. Fig. 1f**; 0.016 ug/g, ∼8-fold reduced from saturation), and closely paralleled the activation of TRGs in cortex (**Supp. Fig. 1a**). These data demonstrate that spontaneous exploratory-like behaviors are highly sensitive to circulating levels of T3.

In contrast, sobetirome administration did not alter the duration mice spent in the exposed area (**Fig. 1e**), despite its peripheral actions that increased energy expenditure levels similar to T3 (**Supp. Fig. 1g**). Finally, chronic treatment (4 weeks) with propylthiouracil (PTU), which interferes with thyroid hormone synthesis and gradually reduces levels of T3^46^, reduced the time mice spent in the illuminated zone (**Fig. 1e**), and decreased TRG expression in cortex (**Supp. Fig. 1h**). Therefore, peripheral actions of thyroid hormone are not sufficient to alter exploratory behavior. Instead, T3 action in the brain directly and bidirectionally alters spontaneous exploratory-like behaviors in mice.

### Single-nucleus sequencing reveals distinct circuit remodeling programs across cell types

To identify the cortical transcriptional programs that may mediate thyroid-dependent changes in exploratory behaviors, we conducted single-nucleus RNA sequencing (snRNAseq) of dorsal frontal cortex, centered on M2, of control mice or mice exposed to T3 for 2.5 days (n = 4 per condition). After quality control filtering (**Supp. Fig. 2**), we analyzed a final dataset of 52,996 cells from T3 animals and 54,746 cells from control animals, which clustered into clear neuronal and non-neuronal cell classes (**Fig. 2a**), and were assigned cell type labels using a recent single-cell motor cortex atlas as reference^47^ (**Supp. Fig. 2**, **Methods**).

**Figure 2.**
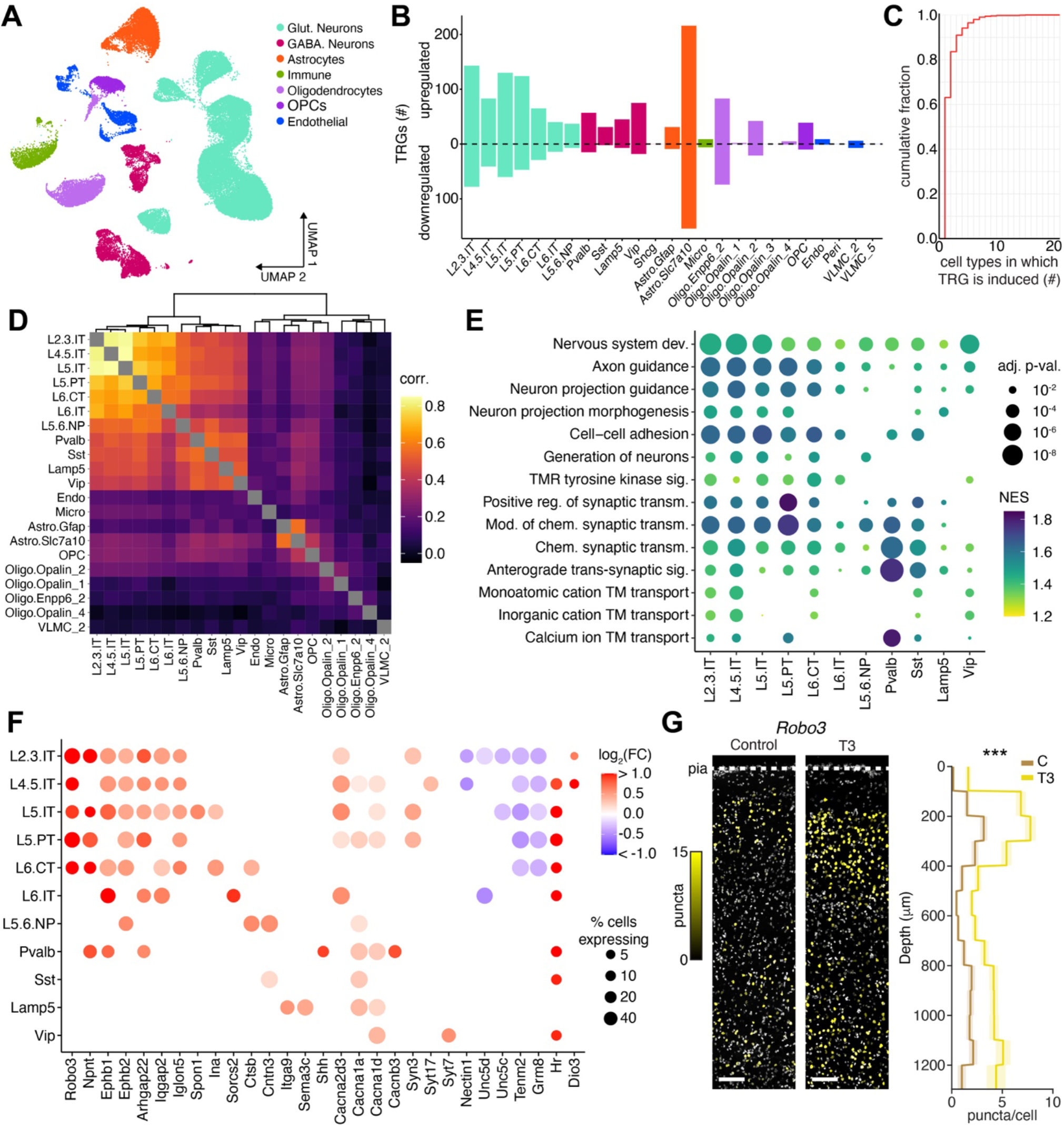
Single-nucleus RNA sequencing of M2 reveals cell type specific T3-induced transcriptional programs. A) UMAP low dimensional representation of nuclear transcriptomes (each dot). Sequenced nuclei readily clustered into broad cell classes (indicated by dot color) and were subsequently mapped onto specific cell types (**Supp. Fig. 2**). B) The numbers of TRGs within each cell type (FDR adjusted p < 0.05, robustness score ≥ 0.5). A full list of identified TRGs can be found in **Supp. Table 1**. C) Cumulative distribution of the number of cell types in which TRGs are induced. Most TRGs (∼60%) were detected in only one cell type. D) Heatmap representation of the spearman correlation of changes in expression of TRGs across cell types. Cell types without TRGs were excluded. The dendrogram shows the hierarchical clustering of cell types based on changes in expression TRGs. E) Dot plot of top terms from gene set enrichment analysis (GSEA) of TRGs in glutamatergic and GABAergic neurons. The union of the top 3 terms per cell type are displayed. The color of each dot indicates the normalized enrichment score for a given term associated with TRGs, and the size indicates the FDR-adjusted p-value. TRGs in both glutamatergic and GABAergic neurons were strongly enriched for synapse associated terms, and, in addition, glutamatergic projection neuron TRGs were highly enriched for axon-guidance associated terms. Terms are abbreviated for convenience, TM: transmembrane, TMR: transmembrane receptor. A full list of GSEA results for each cell type with sufficient sample size for differential expression analysis can be found in **Supp. Table 2**. F) Dot plot of TRGs expression changes across glutamatergic and GABAergic neuron classes. Neuron TRGs were enriched for many genes driving axon-guidance GSEA terms, including *Robo3*, *Npnt*, and ephrins (*Ephb1*, *Ephb2)*— prominently in glutamatergic projection neurons— and genes associated with presynaptic function such as *Cacna2d3*, *Cacna1a*, *Syn3*, and synaptotagmins (*Syt7*, *Syt17*). Downregulated genes included *Grm8*, a presynaptic, putative negative regulator of glutamatergic transmission^115^. The color of the dot indicates the fold-change in expression level, and its size indicates the percent of cells within the given cell type expressing the TRG in the T3 state. G) FISH against *Robo3* across layers of M2 (left: control condition, middle: T3 condition). Scale bars = 100 μm. Right: summary of *Robo3* expression as a function of depth. *Robo3* is upregulated by T3 across cortical layers (***p < 10^-4^, Wilcoxon rank-sum test comparing treatment effect across entire cortical depth; Control: n = 9675 cells; T3: n = 16105; 3 mice per condition). Central line indicates the mean, and shaded area are 95% confidence intervals of the mean.

For each cell type, we identified differentially expressed TRGs between control and elevated T3 conditions (FDR-adjusted p < 0.05, robustness score ≥ 0.5, complete gene lists are in **Supp. Table 1, Methods**). T3 exerted widespread influence on gene expression across most cortical cell types (**Fig. 2b**), with 699 genes showing statistically significant upregulation, and 414 showing statistically significant downregulation. TRGs were largely cell-type specific, with only 16% differentially expressed in three or more cell types (**Fig. 2c**). Hierarchical clustering of TRGs according to their induction across cell types (**Fig. 2d**) revealed distinct TRG programs within neuronal and non-neuronal cell types. Neuronal TRG induction was clustered into two coordinated programs, one specific for glutamatergic projection neurons, and another for GABAergic neurons and the local, non-projecting glutamatergic subtype L5.6.NP. Non-neuronal TRG induction revealed distinct programs for astrocytes, subtypes of oligodendrocytes, and immune and vasculature-associated cell types.

Astrocytes are a major source of T3 in the brain through active transport of the pro-hormone thyroxine (T4) across the blood brain barrier, and local conversion of T4 to T3 by cytoplasmic type 2 deiodinase *Dio2*. Astrocytes are not passive conduits for T3— instead, T3 regulates aspects of astrocyte differentiation, morphogenesis, and maturation^48^. In response to T3, astrocytes downregulated both *Slco1a1*, the major astrocytic transporter of T4, and *Dio2*, revealing a homeostatic mechanism to stabilize T3 levels by reducing both T4 intake from the blood, and T4 conversion to T3 (**Supp. Fig. 3**). We conducted gene set enrichment analysis (GSEA) of astrocyte TRGs to gain insight into the cellular processes influenced by T3 (**Supp. Table 2**). GSEA revealed an over-representation of genes associated with synapse formation and maintenance, including astrocyte expressed genes involved in glutamate release probability and clearance (**Supp. Fig. 3**). These results reveal an astrocyte-specific T3-dependent transcriptional program that may modulate glutamatergic synapses.

Oligodendrocyte progenitor cell (OPC) differentiation and oligodendrocyte maturation are critically dependent on T3^49–52^, and T3 mimetics are being explored as treatments for adult demyelination disorders^53^. Consistent with this, oligodendrocytes and OPCs were responsive to T3, differentially expressing 246 unique TRGs (**Fig. 2b**). GSEA of TRG programs within oligodendrocytes and OPCs (**Supp. Table 2**) identified an enrichment for genes reported as regulators of OPC differentiation and oligodendrocyte myelination, with different programs across OPCs, immature (Oligo.Enpp6_2), and mature oligodendrocytes (**Supp. Fig. 3, Fig. 2d**). Therefore, in response to T3, adult oligodendrocytes and OPCs induce genetic programs— tailored to maturation stage— that likely regulate differentiation, structural remodeling, and myelination.

Within glutamatergic and GABAergic neurons, the top neuronal pathways (**Methods**) from GSEA revealed a striking enrichment for genes associated with axon guidance and plasticity in projecting glutamatergic neurons, and synaptic regulation across both glutamatergic neurons and subtypes of GABAergic neurons (**Fig. 2e**). Consistent with this analysis, induced TRGs included many genes implicated in axon pathfinding, axonal and presynaptic localized cell-cell adhesion, calcium handling, and other presynaptic functions (**Fig. 2f**). Among these genes, *Robo3* stood out as highly induced across most glutamatergic cell types (∼2- to 4-fold increase in expression, **Fig. 2f**), which we confirmed via fluorescence *in situ* hybridization (FISH) across layers of M2 (**Fig. 2g**). Robo3 is a transmembrane protein that, in the context of development, is localized to axons and is required for proper patterning of the nervous system^54^ but whose function in the adult brain has not been explored. Given that changes in transcription do not always lead to changes in protein levels due to post-transcriptional mechanisms, we measured Robo3 protein levels and found a marked increase in mice treated with T3 (∼3-fold, **Supp. Fig. 4**). The distinct TRG programs in neurons elucidated here may contribute to the plasticity of cortical circuits by modifying synaptic connectivity of GABAergic neurons and altering the molecular composition of axons in glutamatergic neurons, thereby inducing downstream processes such as the establishment of axonal fields and glutamatergic synapses. Further, these changes in axon growth or maturation would likely require increased myelination, and the establishment or modification of synapses would likely involve support from astrocytes, consistent with the pro-myelination and synaptic programs induced by T3 in oligodendrocytes and astrocytes, respectively. These transcriptional results suggest that T3 drives a concerted program in multiple cell types to remodel cortical circuits.

### T3 acts locally within glutamatergic projection neurons to alter their transcriptome

In principle, the observed changes in neuronal gene expression in M2 could be a consequence of thyroid hormone action elsewhere in the brain. For example, T3 could activate receptors in the hypothalamus, resulting in secretion of a factor or a change in activity that subsequently alters cortical transcription. Alternatively, neurons may respond indirectly via T3-activated programs within neighboring glia. To address whether the observed T3-regulated transcriptional programs are a cell-autonomous response to T3 in neurons of cortex, we utilized a thyroid receptor β (THRβ) mutant derived from an individual with generalized thyroid resistance^55^. Although the receptor has an intact DNA binding domain, it has a single amino acid deletion (threonine 337, **Fig. 3a**) that prevents it from binding thyroid hormone and activating transcription^56^. Therefore the mutant THR acts as a dominant negative (DN), competing for DNA binding sites, which are largely shared between THRα and THRβ ^57–59^, and blocking transcription in response to T3. Furthermore, unliganded THRs are bound to DNA and recruit transcriptional regulators. This aspect of THR biology is preserved in the DN-THR and would be lost with genetic deletion of thyroid receptors^59^.

**Figure 3.**
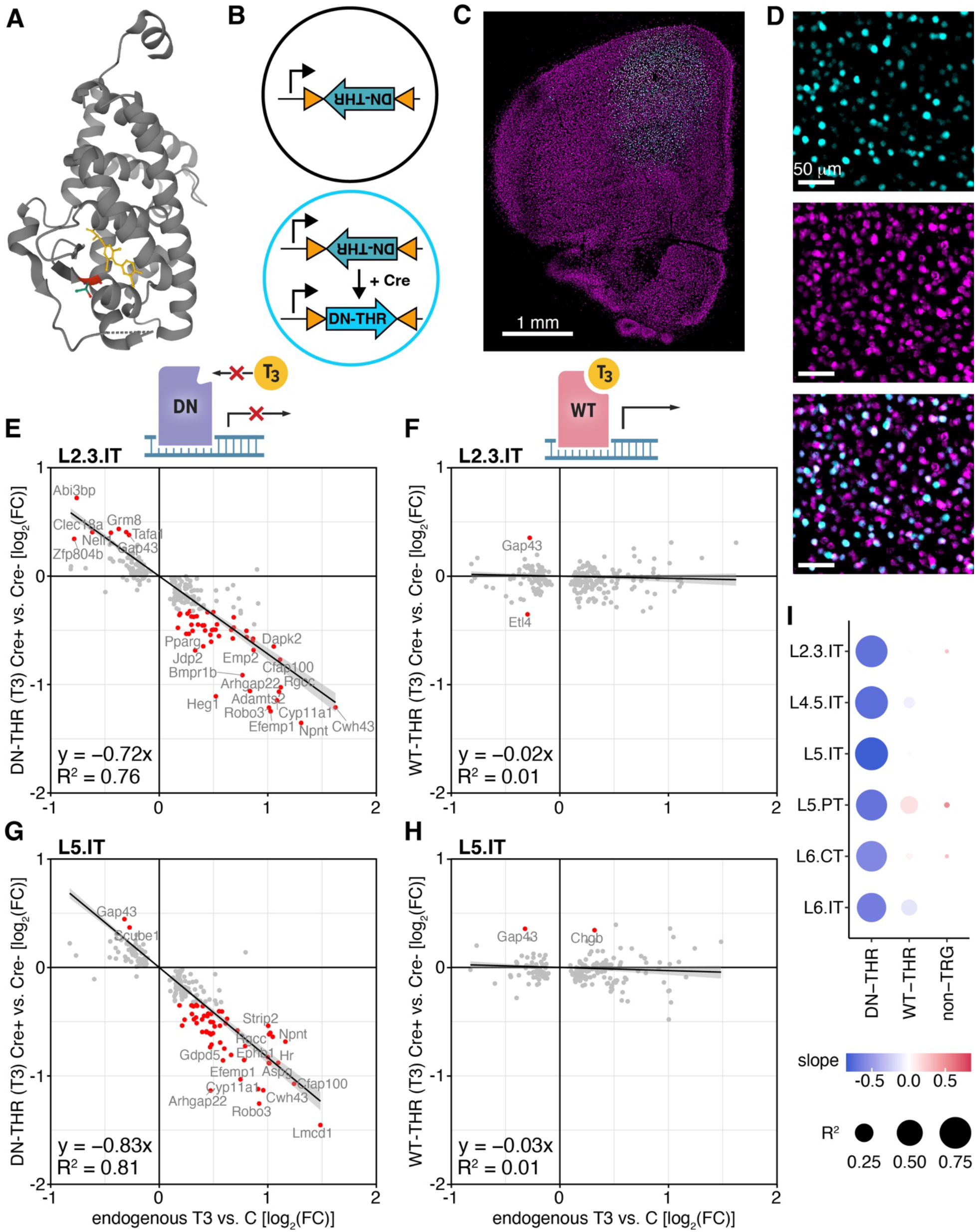
T3-induced transcriptional programs are due to direct T3 action on its receptors. A) Crystal structure of human THR bound to T3 (PDB Entry 3GWS). Threonine 337 is highlighted in red; its deletion in the dominant negative (DN) receptor prevents T3 binding. B) Schematic of viral strategy. An AAV driving expression of a Cre-dependent DN-THR was transduced broadly. A second self-complementary AAV driving expression of Cre was delivered at low infectious titer to produce mosaic tissue in which only a subset of neurons express Cre and activate expression of DN-THR. C) Image of antibody-stained cortical hemisphere showing a representative injection with expression of Cre (cyan) labeled by an HA tag, centered on M2, and all neurons labeled by NeuN (magenta). D) Magnified images within the injection site, showing Cre expressing neurons (top, cyan, HA labeled) and all neurons (middle, magenta, NeuN labeled), and their overlay (bottom). We found Cre expressed in 44% of neurons (1639/3708 NeuN labeled cells were also HA positive, from n = 2 animals). E) Plot showing on the y-axis the log_2_(fold-change) of L2.3.IT TRGs (Fig. 2b) between Cre+ (DN-THR expressing) L2.3.IT cells and Cre-(lacking DN-THR) L2.3.IT cells, after T3 treatment. The x-axis shows the log_2_(fold-change) of L2.3.IT TRGs between the T3 and vehicle control conditions from the original dataset (Fig. 2). Red dots highlight TRGs whose expression was significantly disrupted by 25% or more due to DN-THR (FDR-adjusted p < 0.05, and fractional change in expression of at least ±25%). Linear regression fits to the data are overlaid, grey shading indicates a 95% confidence interval. Fit equation and R^2^ value are displayed on the lower left. See **Supp. Fig. 5** for plots of each cell type. F) As in **E)**, but for WT-THR expressing tissue. The y-axis shows the log_2_(fold-change) of L2.3.IT TRGs between Cre+ (WT-THR expressing) L2.3.IT cells and Cre-(lacking WT-THR) L2.3.IT cells, after T3 treatment. Red dots as in **E)**. See **Supp. Fig. 5** for plots of each cell type. G) As in **E)**, but for L5.IT neurons, and L5.IT TRGs (Fig. 2b). H) As in **F)**, but for L5.IT neurons, and L5.IT TRGs (Fig. 2b). I) Dot plot showing the linear regression fits across glutamatergic projection neurons. DN-THR expression disrupted TRG programs, resulting in a large negative slope across cell types. This indicated that normally upregulated TRGs are downregulated by DN-THR, and normally downregulated TRGs are upregulated by DN-THR. In contrast, slopes were near zero and had low R^2^ for comparisons between WT-THR expressing and lacking cells, indicating that over-expression of the functional receptor does not broadly disrupt TRG programs. Similarly, DN-THR did not disrupt non-TRG expression.

We injected two adeno-associated viral vectors (AAVs) in M2. One drove expression of a Cre-dependent DN-THR. The other was a self-complementary AAV encoding Cre, targeted to neurons via a human synapsin promoter, and delivered at low infectious titer to produce stochastic transduction, yet robust expression of Cre^60^. This viral strategy created mosaic tissue with a subset of neurons expressing DN-THR (**Fig. 3b**, **Methods**). After a 2-week period to allow for receptor expression, we treated animals with T3, and then dissected the tissue and performed snRNAseq. Cre transcripts were detected in ∼44% of neurons, similar to the proportion of Cre positive neurons detected by immunohistochemistry (**Fig. 3c-d**).

We focused on glutamatergic projection neurons, which as a class had the largest sample size and induced the most TRGs among neurons, facilitating statistical analyses. We compared the expression of TRGs in these neuronal cell types with and without DN-THR (defined by the detection of Cre, **Fig. 3eg**, **Methods**). The presence of DN-THR significantly dampened the transcriptional response to T3 across all tested cell types (**Fig. 3eg, Supp. Fig. 5**). For instance, in L2/3 and L5 IT neurons, linear regressions comparing changes in TRGs due to DN-THR versus endogenous TRG induction resulted in large negative slopes driven by DN-THR downregulating the expression of normally upregulated TRGs and upregulating the expression of normally downregulated TRGs (**Fig. 3eg**). These effects were consistent across glutamatergic projection neurons (**Fig. 3egi, Supp. Fig. 5**). Critically, the perturbation caused by DN-THR was selective to TRGs, as non-T3-regulated genes were minimally disturbed (linear regression R^2^ ≤ 0.02 across cell types, **Fig. 3i**), significantly modulating the expression of only 1% of genes by 25% or more (FDR-adjusted p < 0.05, fractional change in expression of at least ±25%, **Supp. Fig. 5**).

To confirm that disrupted TRG expression was due to the inability of DN-THR to bind T3 and activate transcription, and not due to altered composition of thyroid receptors in the cell, we repeated the experiments with a Cre-dependent WT-THR, differing from DN-THR solely by the presence of a single amino acid, threonine 337. In contrast to DN-THR, expression of the WT-THR maintained the TRG program, reflected by linear regression slopes of near zero for all cell types and a lack of significantly disrupted TRGs (**Fig. 3fhi, Supp. Fig. 5**). Collectively, these experiments illustrate that DN-THR expression in individual glutamatergic projection neurons is sufficient to perturb their TRG program. This suggests that the observed neuronal T3-induced transcriptional changes largely result from local T3 binding to, and activation of, its receptors within individual cortical neurons, as opposed to indirect effects mediated by T3 modulations of other brain regions or cell types. Furthermore, these genetically encoded tools can be used to selectively perturb T3-regulated gene expression in specified cell-types and brain regions.

### T3 rewires cortex through changes in synaptic transmission

Given the T3-dependent induction of axon-related genes, we examined if T3 levels affect cortical circuits and synapses. Upper layer intratelencephalic (IT) glutamatergic neurons that project to contralateral M2 are critical to decision making and motor planning and execution^61,62^. We expressed a channelrhodopsin variant^63^, CoChR-GFP, in these IT neurons in one hemisphere of M2 (**Fig. 4a**, **Methods**) through retrograde labeling of their anatomical projections. We then performed whole-cell recordings of L2/3 pyramidal neurons in contralateral M2 within the CoChR-GFP labeled axonal field (**Fig. 4ab, Supp. Table 3**) in acute brain slices prepared from T3-treated and control animals. We varied the intensity of blue-light stimulation over two orders-of-magnitude to characterize post-synaptic currents (PSCs) across the full range of optogenetic stimulation (**Fig. 4c**). The resulting data for each neuron was fit to a sigmoid to obtain a response-profile relating light power to the magnitude of evoked currents.

**Figure 4.**
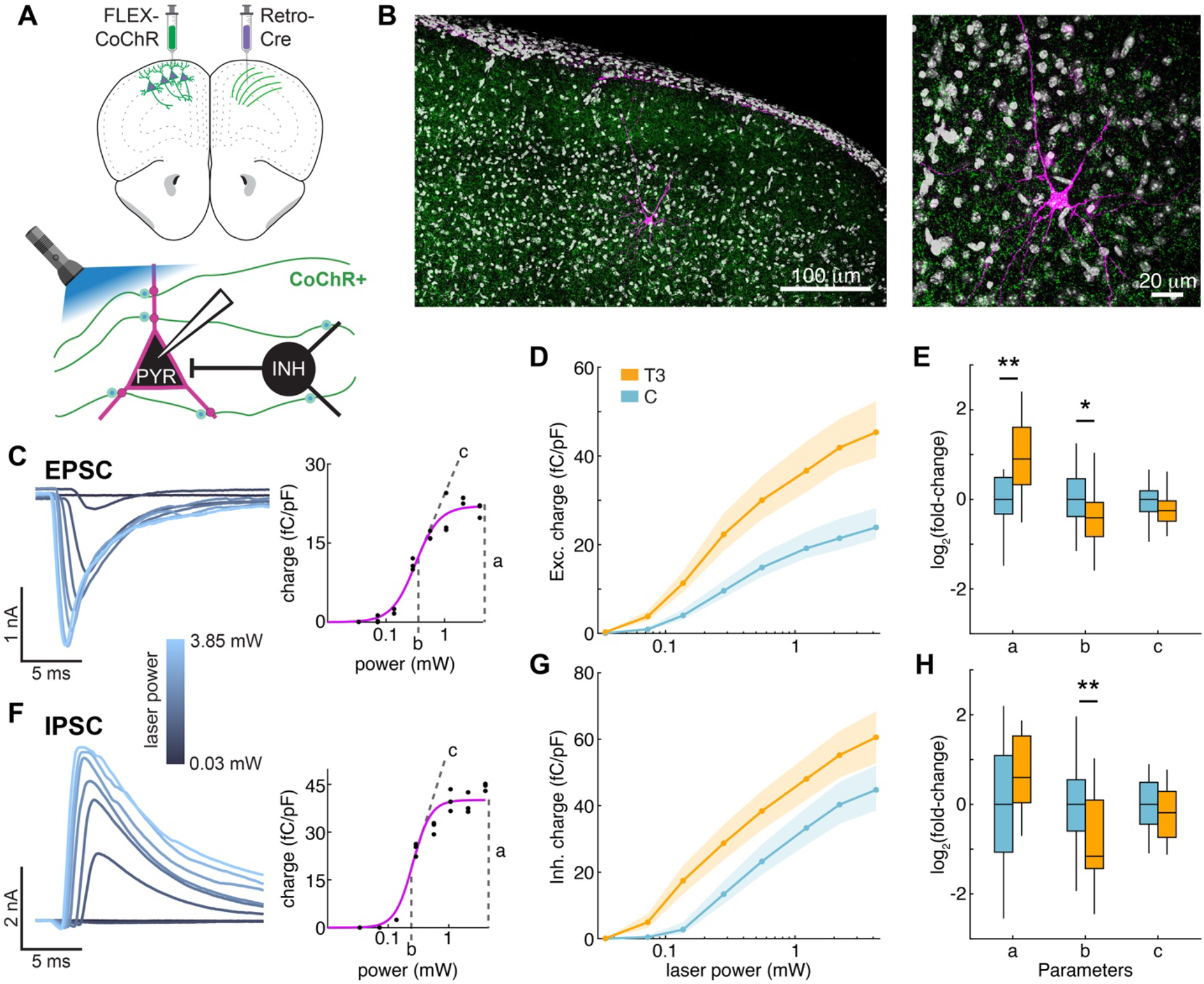
T3 alters synaptic connectivity of cortical glutamatergic neurons. A) Experimental schematic. AAVs encoding Cre-dependent CoChR (FLEX-CoChR) was delivered to the upper layers of M2 in one hemisphere and a retrograde AAV encoding Cre was delivered to the contralateral hemisphere, resulting in CoChR expression in neurons sending projections to contralateral M2 (top). Acute brain slices were prepared and whole-cell recordings were obtained from layer 2/3 (L2/3) pyramidal neurons within the field of CoChR-expressing axons from contralateral M2 (bottom). Post-synaptic currents— both monosynaptic excitatory currents and di-synaptic inhibitory currents—triggered by whole-field light stimulation across a range of intensities were recorded. B) Low magnification (left) and enlarged (right) images of a streptavidin-labeled L2/3 pyramidal neuron filled with biocytin via the patch pipette in a field of CoChR-expressing axons. C) Left, Example of light-evoked excitatory post-synaptic currents (EPSCs) recorded from a recipient L2/3 pyramidal neuron. Currents are color-coded by the light stimulus intensity. Right, Normalized excitatory charge (10 ms window post-stimulus, normalized by cell capacitance) as a function of light stimulus power. Data were fit by a sigmoid characterized by a saturation amplitude (a), a half-maximum inflection point (b, measure of sensitivity), and the slope of the response curve (c, hill coefficient). D) Average normalized post-synaptic excitatory charge as a function of laser stimulus power for T3 (orange, n = 21 neurons, 8 mice) and vehicle (blue, n = 29 neurons, 10 mice) treated mice. Dots represent mean values; shading indicates bootstrapped standard error of the mean. E) Boxplot of changes in each sigmoid parameter (from single cell fits of excitatory charges vs. laser power curves) relative to the median control value. The saturation amplitude (a) was significantly increased by T3-treatment (p = 0.001), and the power to half-maximum (b) was significantly decreased by T3-treatment (p = 0.04). F-H) As in panels **C)-E)**, but for light-evoked inhibitory post-synaptic currents (IPSCs) and normalized inhibitory charge. The power to half-maximum IPSC charge was significantly decreased by T3-treatment (p = 0.005). All statistical comparisons performed with the Mann Whitney U test. * p < 0.05, ** p < 0.01.

T3 treatment increased excitatory post-synaptic currents (EPSCs) across the full range of light intensities (**Fig. 4d**). By parameterizing these responses (**Fig. 4e**), we found that T3-treatment significantly increased the amplitude of saturating post-synaptic responses (p = 0.001) and sensitized the current to light power, reducing the light required for half-maximum stimulation (p = 0.04). In contrast, di-synaptic inhibitory currents (**Fig. 4fg**) onto recipient L2/3 neurons had a significant increase in sensitization (**Fig. 4h**, p = 0.005), but no increase in saturating amplitude (**Fig. 4h**, p = 0.09) in response to T3. The changes in EPSCs were unlikely to be due to alterations in intrinsic excitability of presynaptic neurons, as T3 treatment affected neither the basal properties of L2/3 pyramidal neurons such as membrane capacitance and resistance (**Supp. Table 3**) nor their excitability (**Supp. Fig. 6**). Thus, increasing T3 levels alters synapses to modulate cortical circuits.

To determine whether these synaptic changes are mediated by T3-dependent transcriptional cascades specifically in the pre-synaptic neurons, we co-expressed CoCHR-GFP along with WT-THR (**Supp. Fig. 7a**) or DN-THR (**Supp. Fig. 7f**) in the same IT neurons that send projections to contralateral M2 and repeated the electrophysiology analysis of PSCs in recipient L2/3 pyramidal neurons. The previously observed effects of T3 were recapitulated in experiments with WT-THR expressing in the pre-synaptic neurons (**Supp. Fig. 7a-e**), consistent with the lack of effects of WT-THR expression on T3-dependent transcription (**Fig. 3fhi, Supp. Fig 5**). However, expression of DN-THR in the pre-synaptic neurons occluded the effects of T3 (**Supp. Fig. 7f-j**), suggesting that intact pre-synaptic, T3-dependent transcriptional programs are required to observe T3-induced changes in connectivity.

Finally, we performed similar experiments in *Robo3^fl/fl^* mice, introducing Cre or an inactive Cre control (ΔCre, **Supp. Fig. 8a-e**) into one hemisphere along with CoChR-GFP, and then repeated the electrophysiology analysis of elicited PSCs in recipient L2/3 pyramidal neurons in the contralateral hemisphere. Experiments performed with ΔCre left *Robo3* expression intact, and reproduced PSC changes with T3 (**Supp. Fig. 8a-e**). Loss of presynaptic *Robo3* occluded T3-dependent changes to both direct EPSCs and di-synaptic IPSCs (**Supp. Fig. 8f-j**), indicating that expression changes of *Robo3* in glutamatergic neurons may play a critical role in mediating T3-induced pre-synaptic remodeling of cortical circuits. In sum, these experiments suggest that T3 acts locally in cortical neurons to activate thyroid-dependent transcriptional programs that modify neural connectivity via pre-synaptic mechanisms.

### Effects of T3 on decision making in a probabilistic simulated foraging task

We examined if these thyroid-dependent changes in cortical transcription and circuitry impact exploratory behaviors. Animal foraging is a prototypical exploratory behavior^64^, requiring decisions to be made in an uncertain, dynamic, and seasonal environment. Here, we used a probabilistic simulated foraging task called the 2-armed bandit task (2ABT) which is known, in mice, to require anterior regions of cortex^65–69^. In this task, thirsty, head-restrained mice chose between two spouts located on each side of its head that deliver water with probabilities that change over time (**Fig. 5a**). On each trial, a tone marks the beginning of the selection period, when a mouse can make a choice by licking one of the two spouts. On each trial, one spout, if selected, delivers a drop of water with high probability (0.8), while the other delivers water with low probability (0.2). However, the reward contingencies of the spouts reverse— without any sensory cue— after a block of trials (**Methods**). Therefore, mice must explore both spouts to accumulate evidence about the reward-status of the environment. Mice learned to perform this task and adapt their spout selection in response to the dynamic reward contingencies (**Fig. 5b**).

**Figure 5.**
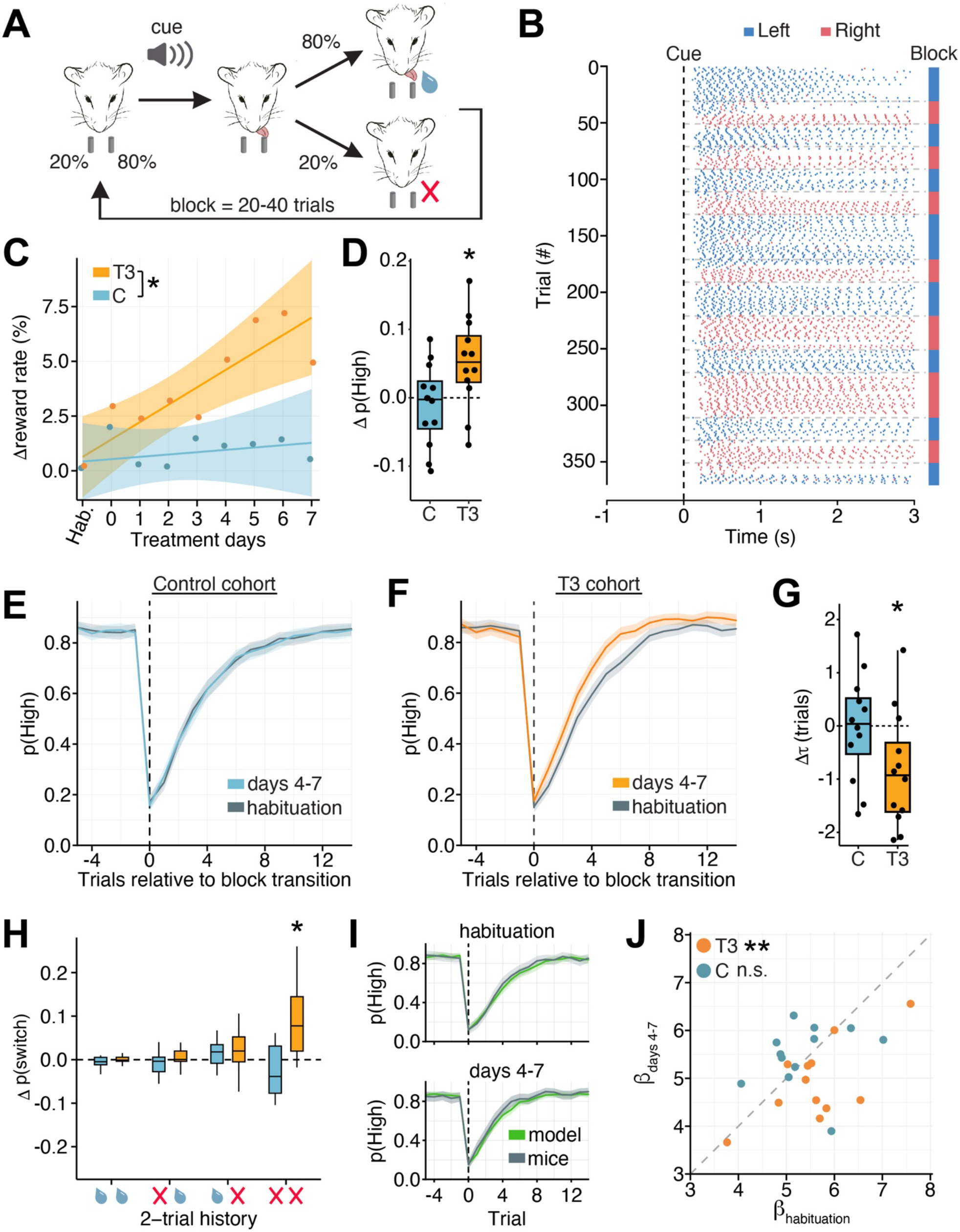
T3 alters decision making and exploration in the 2-armed bandit task (2ABT). A) Schematic of the 2ABT. On each trial, one spout is likely to dispense a water droplet (80%), and the other spout is unlikely (20%). The tone (5 kHz) marks the start of the selection period, during which a mouse can make a choice by licking one of the two spouts. The mouse then receives water drops according to the current reward probabilities of the spouts and its spout choice. These probabilities are dynamic, switching without cue after a block of 20-40 trials. B) Raster plot from a single 2ABT session showing individual licks to left (blue) and right (red) spouts as a function of time from the start of the tone marking the selection period (black dotted line). The first lick in response to the cue is the selection lick and the color code on the right indicates the identity of the highly rewarded spout. C) The percent change in reward rate (rewards/trial) relative to habituation days (Hab.) for T3 (orange) and vehicle (blue) treated mice. The reward rate of each mouse was normalized to the median rate during habituation days. Dots indicate the average change in reward rate per day across mice. Lines are linear fits to the data, and shaded areas are 95% confidence intervals. There was a significant interaction of treatment condition (T3/C) and duration (linear mixed effects model, p = 0.03, likelihood ratio test). D) Change in probability of selecting the highly rewarding spout (p(High)) between the habituation period and days 4-7 of treatment calculated as the difference of median values from each period. Black dots represent single mice. T3 treated mice significantly increased their probability of selecting the highly rewarding spout (p = 0.02, t-test) whereas vehicle treated mice did not (p = 0.55, t-test). E) p(High) as a function of trial position within a block for vehicle treated mice during habituation days (grey) or treatment days 4-7 (blue and orange). Trial 0 marks the block transition (i.e., the first trial of a new block). Shading indicates 95% confidence intervals. F) As in **E)** but for T3 treated mice. G) Change in the time constant (τ) from exponential fits to p(High) after the block transition between the habituation period and days 4-7. Black dots represent single mice. T3 treated mice had a significant decline in τ (p = 0.02, t-test) whereas vehicle treated mice did not (p = 0.93, t-test). H) Change in conditional switch probabilities, dependent on reward outcomes of the previous 2 trials, between the habituation period and days 4-7. The 4 most common histories are plotted, which resulted from selecting the same spout on two consecutive trials with varying reward outcomes, represented by a water droplet (reward) or red X (no reward). T3 mice increased their probability of switching spouts in response to two consecutive failures (p = 0.02, t-test with Benjamini-Hochberg correction). No other conditional switch probabilities changed (p > 0.05, t-test with Benjamini-Hochberg correction). I) Q-learning model predictions of p(High) around block transitions from the habituation period (top) and days 4-7 (bottom). Grey line is the mean probability from the mouse data (the T3 cohort), and the green line is the model prediction. Shaded areas indicate 95% confidence intervals. J) Scatter plot of β parameter fits during habituation (x-axis) and days 4-7 of treatment (y-axis) for each animal. T3-treated mice had a significant decrease in β between habituation and days 4-7 (p = 0.008, t-test), whereas vehicle treated mice did not (p = 0.69, t-test). For all analyses, n = 12 animals for each treatment condition (T3 or control). * p < 0.05, ** p < 0.01.

To determine whether thyroid hormone status influences exploratory decision-making, we trained animals to proficiency, then treated animals with vehicle solution for multiple sessions (habituation period, **Methods**) to establish a baseline behavior, and finally treated animals with T3 (n = 12 mice) or continued vehicle administration (control; n = 12 mice). Motor action measures, such as spontaneous lick rate and reaction times, were not altered by T3 (**Supp. Fig. 9**). However, mouse performance, as measured by the fraction of rewarded trials (reward rate, defined as average reward/trial), improved with T3 (**Fig. 5c**). The reward rate increased with T3 over the 8 days of treatment (p = 0.03, likelihood ratio test; linear regression, F= 12.42 (1,134), p < 10^-3^), consistent with a transcriptional rather than an acute mechanism. In contrast, performance was stable in the control cohort (linear regression, F < 10^-3^ (1,139), p = 0.62, **Fig. 5c**). The T3-induced increase in reward rate was driven by an increase in the probability of choosing the highly reward spout, p(high), which was apparent when comparing performance during the habituation period with that during days 4-7 of treatment (**Fig. 5d**). To understand what changes in behavior underlaid the improvement in task performance, we examined p(high) as a function of trial position within a block (**Fig. 5ef**). With treatment, mice with elevated T3 switched their selection more rapidly to the new highly rewarding spout after block transitions, reflected by a ∼30% smaller time constant from exponential fits of the data (p = 0.02, **Fig. 5g**).

To perform this task optimally, the mice must infer which spout is highly rewarding based on the information collected during previous trials and then use this information to select a spout to sample on the next trial. The faster switch (i.e., in fewer trials) to the new highly rewarding spout after the block transition suggests that animals are not simply switching randomly, but instead altering how they integrate information across trials to infer the highly rewarded spout. To investigate this possibility, we examined the probability of switching spouts after the 4 most highly occurring 2-trial histories (> 50 times per mouse) and compared these conditional switch probabilities during habituation and days 4-7 of treatment. Each 2-trial history resulted from licking the same spout in two consecutive trials with varying reward outcomes (i.e., reward→reward, no reward→reward, reward→no reward, and no reward→no reward) (**Fig. 5h**). T3 treated mice significantly increased the spout switch rate in response to two consecutive failures to receive a water droplet (p = 0.02), while not changing their switching rates in response to the other 2-trial histories. Thus, T3 promotes spout switching dependent on previous choice outcomes that imply a change in reward contingencies.

This paradigm requires mice to engage in two separable processes. First, mice must accumulate evidence to make inferences about the reward-status of the environment. Second, mice use a policy to choose a spout based on the inference. To understand the computations that may govern the observed changes in history-dependent decision making with T3, we modeled the data using a reinforcement learning framework used to understand the separable inference and policy processes in the context of the 2ABT (Q-learning, **Supp. Fig. 10**)^66,70,71^, and whose variables are represented in the firing activity of neurons in anterior regions of cortex during the task^66^. The model fit the data well, both prior to and during T3 treatment (**Fig. 5i**, spout-choice prediction accuracy on held-out data during habituation: 0.85 ± 0.03, mean ± std. dev.; and during days 4-7 of treatment: 0.85 ± 0.03; comparison between epochs: p = 0.52). The model parameters include learning (α) and forgetting (ζ) rates for chosen and unchosen spouts, neither of which changed with T3 treatment (**Supp. Fig. 10**), suggesting that T3 did not alter rates of evidence accumulation or storage. The only parameter to change with T3 was β, which characterizes the animal’s policy by determining the degree to which decisions exploit current evidence relating to which spout is highly rewarding versus exploring the spouts stochastically to gain new information. β decreased with T3 (**Fig. 5j**, p = 0.008), enabling animals to switch spouts more rapidly at block transitions, and improve their overall task performance. Collectively, these results show that mice with increased T3 levels selectively change their decision-making policy to explore the alternate spout with less evidence that the environment has changed.

Finally, we asked whether the observed changes in behavior depended on the neuronal-specific TRG program in anterior regions of cortex. After training animals to proficiency, we introduced AAVs driving expression of DN-THR or WT-THR in neurons centered on M2 in frontal cortex (**Fig. 6ab, Supp. Fig. 11-12, Supp. Table 4**). We continued training the animals for ∼2 weeks to allow for THR expression. We then carried out the same paradigm as before— a habituation period followed by 8 days of T3 treatment. T3 did not induce alterations in gross motor actions such as spontaneous lick rate or reaction times in either the WT-THR cohort (n = 13 animals) or the DN-THR cohort (n = 14 animals) (**Supp. Fig. 13**). The WT-THR cohort recapitulated the previous observations of T3 effects on performance: mice increased their reward rate over treatment sessions (**Fig. 6c**, linear regression, F= 14.14 (1,154), p < 10^-3^), mediated by an increased probability of selecting the highly rewarding spout (**Fig. 6d**, p = 10^-3^). Animals more rapidly switched their selections after block transitions with T3 (**Fig. 6e, g**), and their probability of switching was altered dependent on previous trial outcomes (**Fig. 6h**, T3 significantly increased switching after two consecutive failures, p < 10^-3^). Finally, when their behavior was modeled, β declined with T3 (**Fig. 6i,** p = 0.007), while all other parameters remained unchanged (**Supp. Fig. 13**).

**Figure 6.**
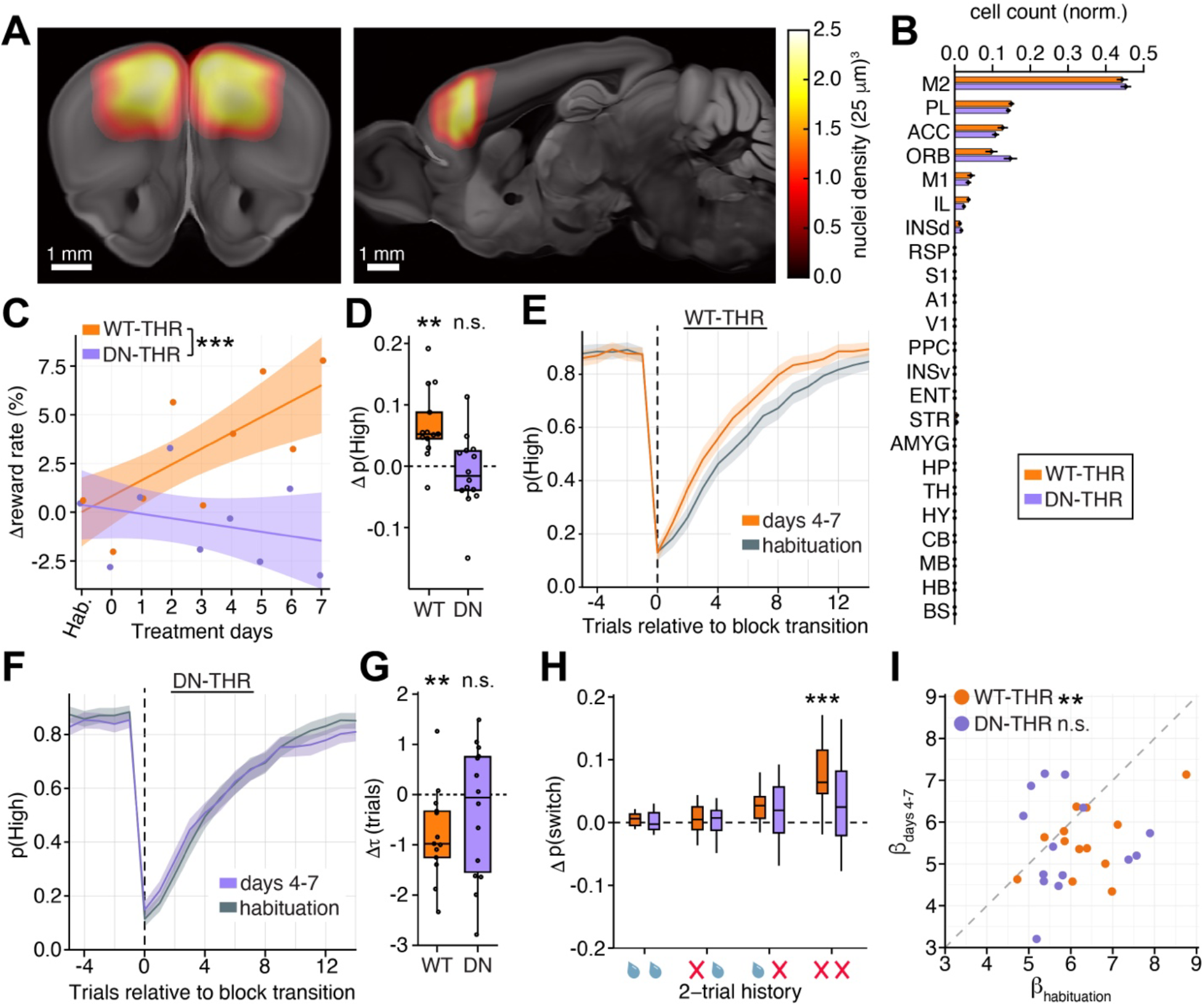
T3-dependent transcriptional cascades in frontal cortex underlie T3 mediated changes in exploratory decision making. A) Average heatmaps of DN-THR expression in the DN-THR cohort of mice performing the 2ABT, registered and overlaid on the Allen Mouse Brain Common Coordinate Framework average template. AAVs encoding WT- or DN-THR were delivered via intracranial injections, and expression was driven by a hSyn promoter for selective expression in neurons (**Supp. Fig. 11**). Scale bar: 1 mm. The coronal section is at ∼2 mm anterior to bregma, and the sagittal section is ∼1 mm from the midline. An extended selection of heatmaps of both DN-THR and WT-THR animals are in **Supp. Fig. 12**. B) Bar plot of the number of WT-THR (orange) and DN-THR (purple) positive nuclei across brain regions. There were no significant differences in brain region expression between the WT-THR and DN-THR cohorts (see **Supp. Table 4** for full details). C) The percent change in reward rate (rewards/trial) over habituation and treatment. All habituation days are grouped (Hab.), and the reward rate of each mouse was normalized to the median rate during habituation. Dots indicate the average change in reward rate per day for WT-THR (orange) and DN-THR (purple) animals. Both cohorts received T3. Lines are linear fits to the data, and shaded areas are 95% confidence intervals. There was a significant interaction of genotype (WT/DN) and treatment duration (linear mixed effects model, p < 10^-3^, likelihood ratio test). D) Change in p(High) between the habituation period and days 4-7 of treatment calculated as the difference of median values from each period. Black dots represent single mice. WT-THR mice significantly increased p(High) (p = 0.001, t-test) while DN-THR mice did not (p = 0.54, t-test). E) p(High) for WT-THR animals as a function of trial position within a block. Orange line indicates mean probabilities over days 4-7. Grey line indicates mean probabilities over habituation days. Shading indicates 95% confidence intervals. F) As in panel **E)** but for DN-THR animals. The purple line indicates mean probabilities over days 4-7, the grey line indicates mean probabilities over habituation days. Shading indicates 95% confidence intervals. G) Change in the time constant (1) of recovery of p(High) from exponential fits of the data (aligned to the block transition) between the habituation period and days 4-7. Black dots represent single mice. WT-THR mice had a significant decline in 1 (p = 0.009, t-test) whereas DN-THR mice did not (p = 0.35, t-test). H) Change in conditional switch probabilities, dependent on reward outcomes of the previous 2 trials, between the habituation period and days 4-7. The 4 most common histories are plotted, which resulted from selecting the same spout on two consecutive trials with varying reward outcomes, represented by a water droplet (reward) or red X (no reward). WT-THR mice increased their probability of switching spouts in response to two consecutive failures (p < 10^-3^, t-test with Benjamini-Hochberg correction). No other conditional switch probabilities changed (p > 0.05, t-test with Benjamini-Hochberg correction). I) Scatter plot of β parameter fits during habituation (x-axis) and days 4-7 of treatment (y-axis) for each animal. WT-THR mice had a significant decrease in β between habituation and days 4-7 (p = 0.007, t-test), while DN-THR mice did not (p = 0.28, t-test). For post-hoc histology and quantification in **A)** and **B)**, WT-THR cohort: n = 12 animals, DN-THR cohort: n = 10 animals. For all other analyses, WT-THR cohort: n = 13 animals, DN-THR cohort: n = 14 animals. ** p < 0.01, *** p < 10^-3^.

In contrast, DN-THR expressing animals no longer responded with a marked change in performance in response to elevated T3. The DN-THR cohort had no increase in reward rate over the experimental time course (**Fig. 6c**, linear regression, F= 1.11 (1,166), p = 0.29), did not change their probability of selecting the highly rewarding spout (**Fig. 6d**, p = 0.54), and did not alter their trial-outcome dependent switching probabilities (**Fig. 6f-h**). In addition, there were no significant changes in any Q-learning modeling parameters (**Supp. Fig. 13**), including β (**Fig. 6i**, p = 0.28) after T3 administration. Thus, although thyroid receptors are expressed in many organs and in many brain areas, notably the hypothalamus, T3-dependent transcriptional cascades in neurons within frontal cortex are necessary for thyroid mediated adaptive changes in exploratory decision making. These studies reveal a mechanism by which a hormonal sentinel of environmental change directly affects neuronal circuits in cortex to drive adaptive changes in exploratory drive.

## Discussion

Variation in thyroid hormone levels can have profound effects on behavior ranging from depressed-like states with pathologically reduced thyroid function to manic or even psychotic states with pathologically elevated thyroid hormone in humans with hypo- and hyper-thyroidism respectively^29–32^. Similar behavioral changes occur in wild and domesticated animals with seasonal changes in thyroid function, leading to adaptive changes in exploratory behavior in synchrony with environmental variables such as food availability and weather. Here we examine the mechanisms of such changes and reveal a T3-sensitive transcriptional program induced in mouse cortical neurons that rewires cortical circuits and increases exploration and risk taking in a variety of contexts. Our findings reveal that cortical axon pathfinding transcriptional programs can be reawakened and are dynamically modulated in adult cortex and serve a physiological function to link peripheral organ metabolism to central exploratory drive. Pharmacological manipulation of such programs may have therapeutic value for the treatment of neuropsychiatric disorders such as depression and bipolar disorder in which exploratory drive is dysregulated.

### T3-induced cell-type specific transcriptional programs

Mammalian cerebral cortex contains dozens of transcriptionally defined cell types, yet the transcriptional signatures that demarcate these cell types are not static. Instead, they change over development and aging^72,73^, and respond dynamically to life experiences^74,75^. Likewise, hormones fluctuate in response to experience, environment, age, and internal somatic states over timescales ranging from minutes to seasons to lifespans^76–78^. Nuclear hormone receptors, which are often ligand-dependent transcription factors^79^, are widely expressed in cortex, and cortical cell types and circuits are responsive to a wide array of hormonal factors^5,80–82^. Therefore, circulating hormones are likely prominent contributors to the dynamic transcriptional landscape of cortex. Here we have characterized one such signaling cascade and elucidated its role in shaping animal behavior.

We find that thyroid hormone levels, known to regulate the metabolic state of peripheral organs, induce cell-type specific transcriptional programs in adult cerebral cortex, and, at least for glutamatergic projection neurons, this induction is locally driven by T3 and is largely cell-autonomous. These programs are tailored to the function of each cell type. For instance, glutamatergic projection neurons engage programs highly enriched for molecules involved in axonal remodeling, while both astrocytes and neurons are enriched for molecules involved in assembling and regulating synapses. In oligodendrocytes, T3 regulated pathways related to differentiation, maturation, and myelination. Although the T3-dependent program in neurons is necessary for T3-dependent plasticity of cortical circuits and subsequent changes to exploratory behaviors, T3 induced programs across all cell types, including glia, likely act in concert to sculpt T3-responsive adaptive behaviors. Future studies are necessary to determine the roles that each cell-type specific program plays in remodeling cortex and behavioral outputs.

### Hormonally induced remodeling of adult cortical circuits

Although synaptic plasticity occurs in adult cerebral cortex and is thought to be essential for learning, many aspects of adult cortical circuits appear largely static. Axons, dendrites, and the size and placement of dendritic spines of glutamatergic neurons are relatively stable, especially when compared to juvenile brains, over periods of weeks and months^83^. However, structural plasticity can be induced in adult cortex with experience, learning, local or even peripheral injury^84–87^. Although most structural plasticity studies have focused on dendrites and dendritic spines, axons and large-scale white matter tracts can also remodel in adult cortex^87–92^.

We find that increased T3 induces expression of genes in cortical glutamatergic neurons that contribute to proper patterning of the nervous system during development, suggesting modulation of circuit patterning programs. Consistent with their axon pathfinding function in development, the T3-induced transcriptional program in glutamatergic neurons drives plasticity of cortical circuits via pre-synaptic mechanisms. These changes manifest as alterations in the magnitude of cortical excitation and recruitment of polysynaptic inhibition. The structural substrate of this plasticity remains to be elucidated and, consistent with the function of the induced genes, could include axon growth and branching with new synapses or synapse formation on existing axons. Indeed, T3 induces expression of synaptic genes in both glutamatergic and GABAergic neurons, which can contribute to the reported effects independent of axonal remodeling. In addition, given the induction of pro-myelination genes by T3 in oligodendrocytes, and the activation of synaptic regulators in astrocytes, non-neuronal cells likely contribute to circuit remodeling. A full understanding of the role each of these axon-guidance and synapse associated molecules plays in T3-induced cortical circuit plasticity will reveal the molecular and anatomical mechanisms driving this novel form of hormonally induced adult plasticity in cerebral cortex.

### Coordination of exploration and metabolism across species

We find that T3-driven transcriptional programs in neurons of anterior cortex causally drive changes in exploratory drive, such that higher levels of thyroid hormone synchronize an increase in body-wide metabolism with an increased tendency towards exploratory, information-seeking behaviors. Changes in exploratory behaviors occurred across paradigms, including coarse paradigms of exploration such as the light-dark (LD) preference assay, as well as in theoretically motivated and learned paradigms, such as the 2ABT, designed to probe exploratory decision-making. Importantly, the LD assay, which measures an innate aversion of mice to well-lit, exposed environments, shows bidirectional modulation with changes in thyroid function, suggesting that basal T3 levels modulate the propensity of the mouse to overcome this aversion. Although the motivating drive to explore a perceived risky environment is likely multifaceted, the result can be interpreted as a change in the tradeoff between information gained versus evaluated risk, fundamental to exploratory behaviors. Coupled with our findings from the 2ABT, which revealed a specific increase in information-seeking decision-making, the LD preference assay can serve as a first-pass behavioral screen for T3-dependent molecular modifiers of exploratory behavior.

The robust, coordinated response of body and brain to thyroid hormone is likely adaptive. In environments with seasonal changes in food availability, an increase in thyroid hormone signals an increase in the expected abundance of resources. The sensory cues promoting changes to thyroid hormone signaling remain to be fully elucidated and may vary between species, but drivers include photoperiod, temperature, and diet^21,93–95^. Regardless of sensory triggers, during the time of year when food is available, animals’ thyroid hormone levels rise, increasing their energy expenditure and willingness to explore and forage. Conversely, during times of year when resources are scarce, thyroid hormone levels drop, slowing metabolism and biasing animals towards energy conserving behaviors. Indeed, thyroid hormone levels are known to drop in response to food restriction and deprivation^96^, and low T3 levels within hypothalamus are likely critical for maintaining a state of hibernation^16,97^.

Evidence in support of adaptive changes in thyroid hormone levels is prevalent in human populations. Seasonal fluctuations in HPT-axis function are ubiquitous^98^, and human populations that experience large changes in temperature and photoperiod between winter and summer months also exhibit large seasonal fluctuations in T3^99,100^, similar to thyroid fluctuations observed in wild animals. Beyond seasonal variation, thyroid levels respond dynamically to life experience. For instance, a wide array of both acute and chronic systemic illnesses sharply depress T3 levels^101^. This response can be considered adaptive in light of our results, as it biases the sick individual towards isolation and energy preserving behaviors. Outside of pathologies, the effects of normal variation of thyroid hormone across humans with no thyroid dysfunction covaries with population-level socio-economic outcomes. Physiological variation in T3 is strongly correlated with the likelihood of being employed and total hours worked in representative US adult populations^102^. Although the relationship to employment status and drive to work is compelling, further studies are necessary to detail the effects of physiological thyroid levels on cognition and exploratory behaviors in humans. Given that thyroid hormone levels can be reliably measured in standard blood tests, and that imaging techniques such as MRI can measure patterns of activity across the human brain during cognition, and diffusion tensor imaging can resolve changes in white matter comprised of axonal tracts and myelin, mechanistic human studies are readily achievable.

### The effects on brain function and behavior due to T3 level decline over the lifespan

Another dynamic feature of thyroid physiology in humans and other species is a steep decline in T3 levels with aging^102^. This change is thought to contribute to reducing metabolic rates in aging populations, but its contribution to age-dependent changes in brain function has yet to be elucidated. In humans and animals, exploratory and risk-taking behaviors peak during adolescence before declining with age^103,104^. Conversely, the prevalence of anhedonia and depression increases with age^105^, consistent with the characterization of depression as a pathological neglect of exploratory behaviors. Intriguingly, T3 has been used as an effective therapy for depression in humans^106^ independent of thyroid status, supporting our results that thyroid hormone levels regulate exploratory drive. In humans, changes in 2ABT performance have been observed between younger and older adults^107^. It will therefore be interesting to investigate whether age-dependent alterations in exploratory behaviors can be quantified in the 2ABT established here, and whether these effects can be ameliorated by increasing T3 concentrations within the brains of old mice.

Furthermore, brain-targeted thyroid mimetics are in development for the treatment of other age-related brain disorders, such as multiple sclerosis^53^, whose degenerating cell type, oligodendrocytes, responds robustly to T3 by inducing a gene set enriched for remyelination and regeneration described above. It is therefore possible that restoring youthful T3 levels in an organ-specific manner could not only boost bodily metabolism, but also combat a range of neuropsychiatric and neurodegenerative diseases whose prevalence increases with age.

### Cortical circuits that contribute to mania

Our findings suggest that changes in thyroid hormone levels are a natural perturbation that modulate cortical circuitry along physiologically relevant axes, resulting in altered exploratory drive. Such evolutionarily conserved programs^108^, including thyroid hormone-dependent control of behavior, can be exploited to reveal canonical brain circuit functions that go awry in psychiatric disorders in which exploratory drive is dysregulated. These include depression (discussed above) and symptoms of mania and risk-taking present in bipolar disorder (BD). The overlapping symptomatic profiles of BD and hyperthyroidism are intriguing. Individuals with BD have increased prevalence of thyroid disease^109^, treatment with lithium is known to significantly perturb thyroid organ function^110^ and thyroid receptor expression in the brain^111^, and thyroid modulators can be effective for BD management^112^. More broadly, there is a high incidence of metabolic disorders, such as diabetes mellitus^113,114^, in individuals with BD, suggesting a perturbed link between exploratory behavior and peripheral metabolism. Despite etiological differences, the challenges of modeling BD using monogenic animal models motivate a search for convergent mechanisms that could exist at the molecular, cellular, and circuit levels between these disorders, providing a path to search for conserved targets for disease treatment.

## Conclusion

Our studies reveal how the action of a single hormone can coordinate two seemingly disparate biological phenomena: exploratory drive and whole-body metabolic state. We anticipate that the systematic characterization of the vast array of hormonally driven molecular programs in cerebral cortex will reveal a range of novel circuit plasticity mechanisms to tune complex behaviors to match the needs of the body and the demands of the environment.

**Supplemental Figure 1.**
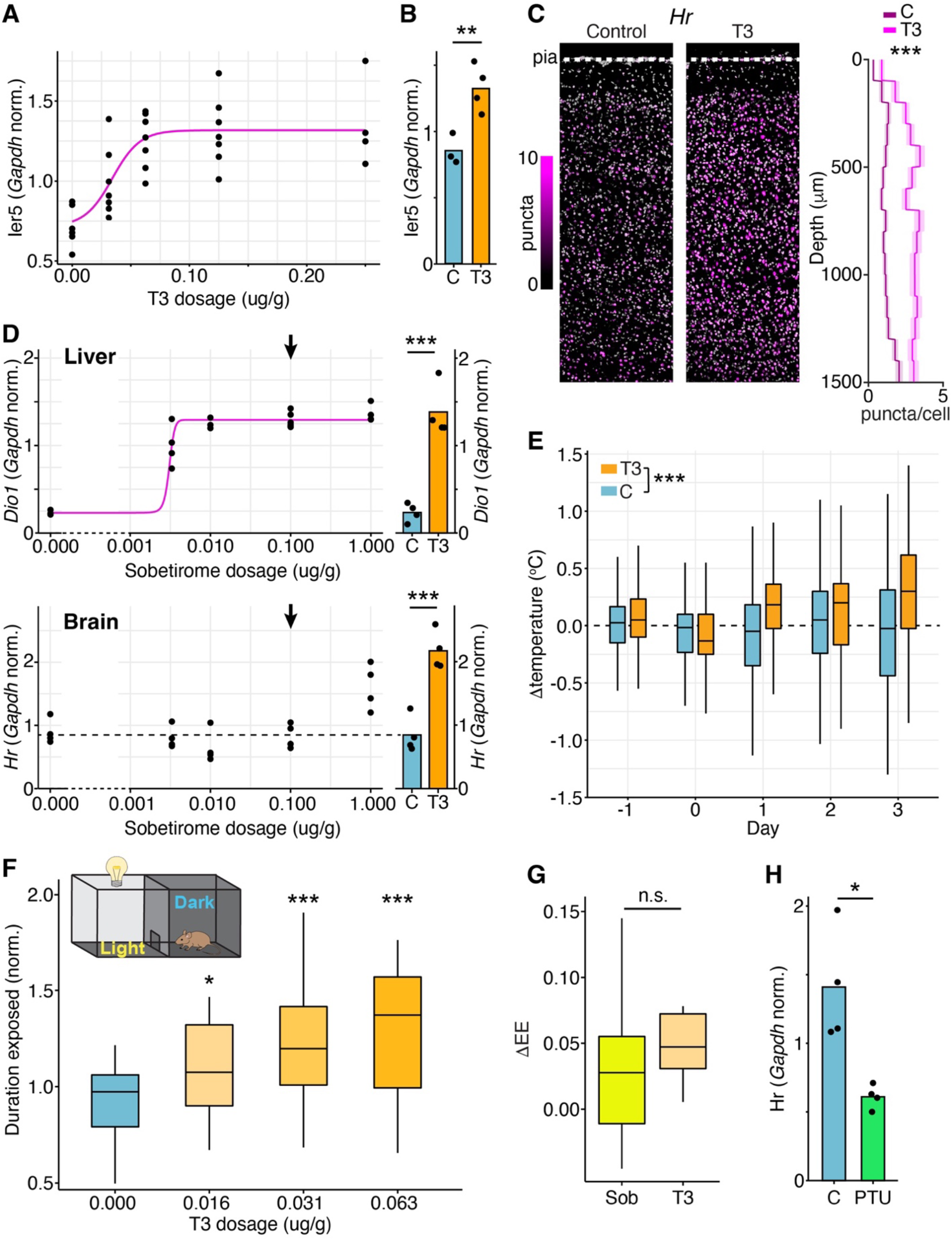
Characterization of T3 effects on cortical transcription, animal physiology, and behavior. A) *Ier5* induction in frontal cortex as a function of T3 concentration. *Ier5* expression saturates by 0.125 ug/g. n = 4-7 mice per condition. Pink line fits the data to a sigmoidal curve. This dose was used for all subsequent experiments unless otherwise noted. A) B) *Ier5* is upregulated in frontal cortex 1 hour after treatment with T3 relative to vehicle control (control: n = 3 mice; T3 treated: n = 4 mice; measured by qPCR; p =0.008 Welch’s t-test). B) FISH of the TRG *Hr* in secondary motor cortex (left: control condition, middle: T3 condition). Scale bars = 200 μm. Right: summary of *Hr* expression as a function of depth. *Hr* is upregulated by T3 across all layers of cortex (***p = 0, Wilcoxon rank-sum test comparing treatment effect across entire cortical depth; Control: n = 5494 cells; T3 treated: n = 6061; 2 mice per condition). Central line indicates mean, shaded area is 95% confidence interval of the mean. C) Top: induction of the TRG *Dio1* in the liver as a function of Sobetirome concentration (left) or with T3 treatment (right). *Dio1* expression differed among treatments (one-way ANOVA, F(6,23)= 45.4, p < 10^-4^). Both T3 (p < 10^-4^, Tukey HSD test) and 0.1 ug/g Sobetirome (indicated by arrow; p < 10^-4^, Tukey HSD test) treatments increased *Dio1* expression in the liver, and were indistinguishable from each other (p = 0.95, Tukey HSD test). Pink line fits the data to a sigmoidal curve. Bottom: induction of the TRG *Hr* in the brain (frontal cortex) as a function of Sobetirome concentration (left) or with T3 treatment (right). *Hr* expression differed among treatments (one-way ANOVA, F(6,23)= 20.0, p < 10^-4^). T3 treatment (p < 10^-4^, Tukey HSD test) increased *Hr* expression in the brain, while 0.1 ug/g Sobetirome did not (indicated by arrow; p = 0.99, Tukey HSD test). 0.1 ug/g Sobetirome was chosen to maximize peripheral TRG induction without central TRG induction. D) Rectal temperature was recorded during habituation (used for baseline temperature) and treatment (Day 0-3). T3-treated animals increased body temperature relative to controls, and the effect of T3 increased over days of treatment (p < 10^-3^, likelihood ratio test of linear mixed effect models; n = 82 mice per condition). E) Duration mice stay exposed in the lit section of the light-dark box as a function of T3 concentration (C: n = 41 mice; 0.016 ug/g T3: n = 32 mice; 0.031 ug/g T3: n = 37 mice; 0.063 ug/g T3: n = 22 mice). Durations from individual experimental replicates were normalized to the median duration of their respective control cohort. There was a significant effect of treatment (χ^2^(3) = 25.423, p < 10^-4^, Kruskal-Wallis’s test), with each T3 concentration resulting in a significant increase in duration exposed relative to control (0.016 ug/g T3: p = 0.01, 0.031 ug/g T3: p < 10^-4^, 0.063 ug/g T3: p < 10^-4^; Dunn’s post-hoc test using Holm’s adjustment for multiple comparisons). F) Sobetirome (0.1 ug/g; n = 17 mice) vs T3 (0.016 ug/g; n = 14 mice) treatment of mice resulted in a change in energy expenditure after 3.5 days of treatment that was similar (p = 0.31, Welch’s t-test). The change in energy expenditure was calculated between treated mice and the average of control cohorts that were run simultaneously (sobetirome vehicle controls: n = 18 mice; T3 vehicle controls: n = 13 mice). G) PTU treatment over 2 weeks led to a downregulation of the TRG transcript *HR* in secondary motor cortex relative to controls (measured by qPCR; n = 4 mice per condition; p = 0.03, Welch’s t-test). For all panels of figure, * = p < 0.05, ** = p < 0.01, *** = p < 0.001. Box plot central lines indicate medians. Box and whiskers represent quartiles 1-4.

**Supplemental Figure 2.**
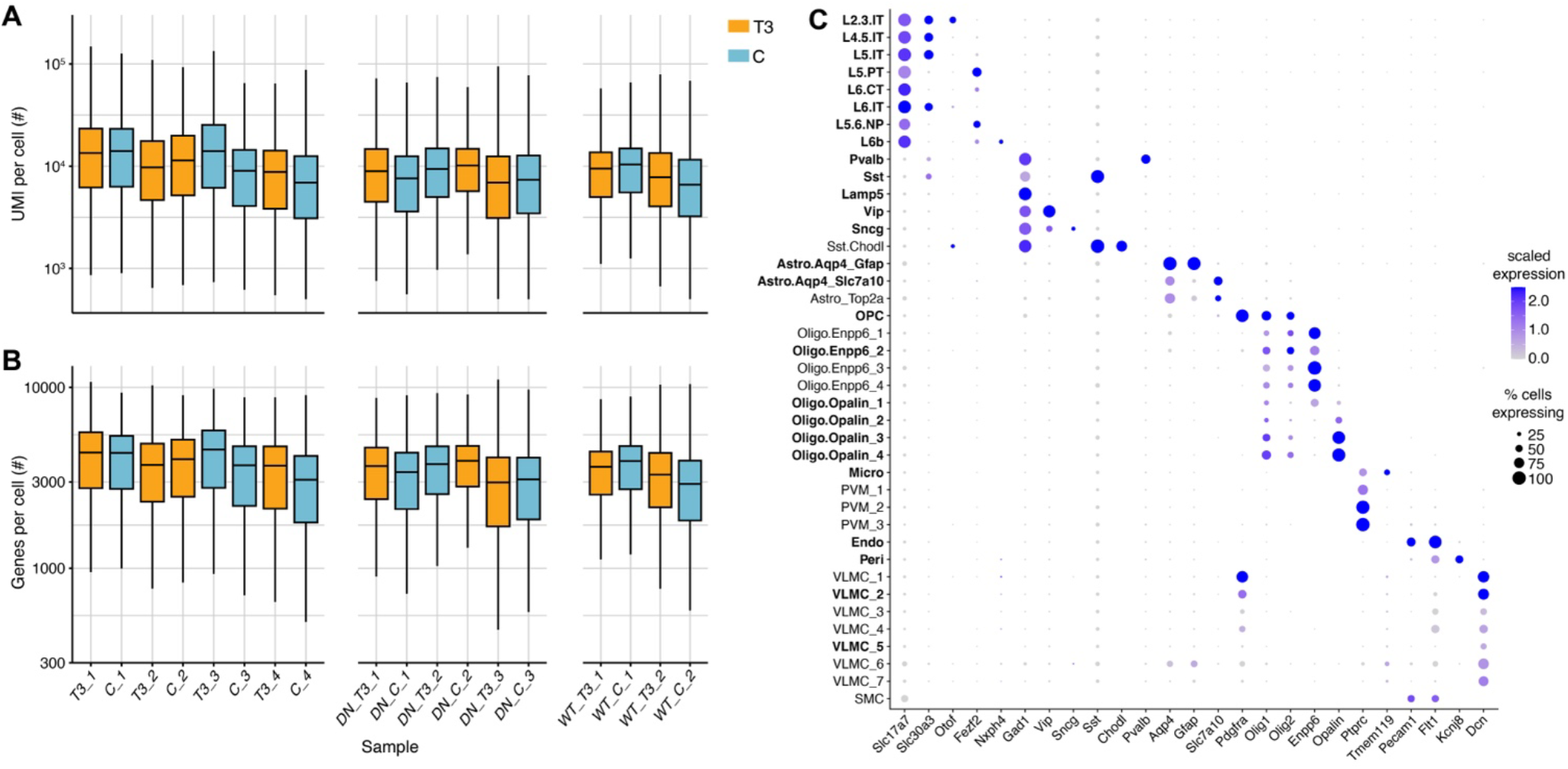
Quality control metrics for single nucleus RNA sequencing. A) Boxplot showing the distribution of the number of unique molecular identifiers (UMIs, putative transcripts) detected in each nucleus over all samples, after quality control filtering. Samples (T3 treated in orange, vehicle treated controls in cyan) showed consistent UMI distributions. Samples T3_1-4 and C_1-4 are from C57BL6/J mice treated with either T3 or vehicle control solutions and reported in **Fig. 2**. Samples DN_T3_1-3, DN_C_1-3, WT_T3_1-2, WT_C_1-2, refer to animals that received intracranial injections of AAVs encoding Cre along with a Cre-dependent dominant-negative (DN) or wildtype (WT) thyroid receptor, and treatment with T3 or vehicle. A subset of these samples (DN-THR, T3: 47,876 cells, WT-THR, T3: 45,076 cells) are reported in **Fig. 3**. B) As in **A)** but for the number of genes expressed in each cell. Samples showed consistent gene expression distributions. C) Marker gene dot plot. After clustering into broad cell classes, cells were mapped onto previously defined cell types from a single-cell atlas of motor cortex^47^. Mapped cell types displayed appropriate expression of marker genes. Cell types in bold had sufficient cell counts to perform analysis of differential gene expression between T3 and C conditions.

**Supplemental Figure 3.**
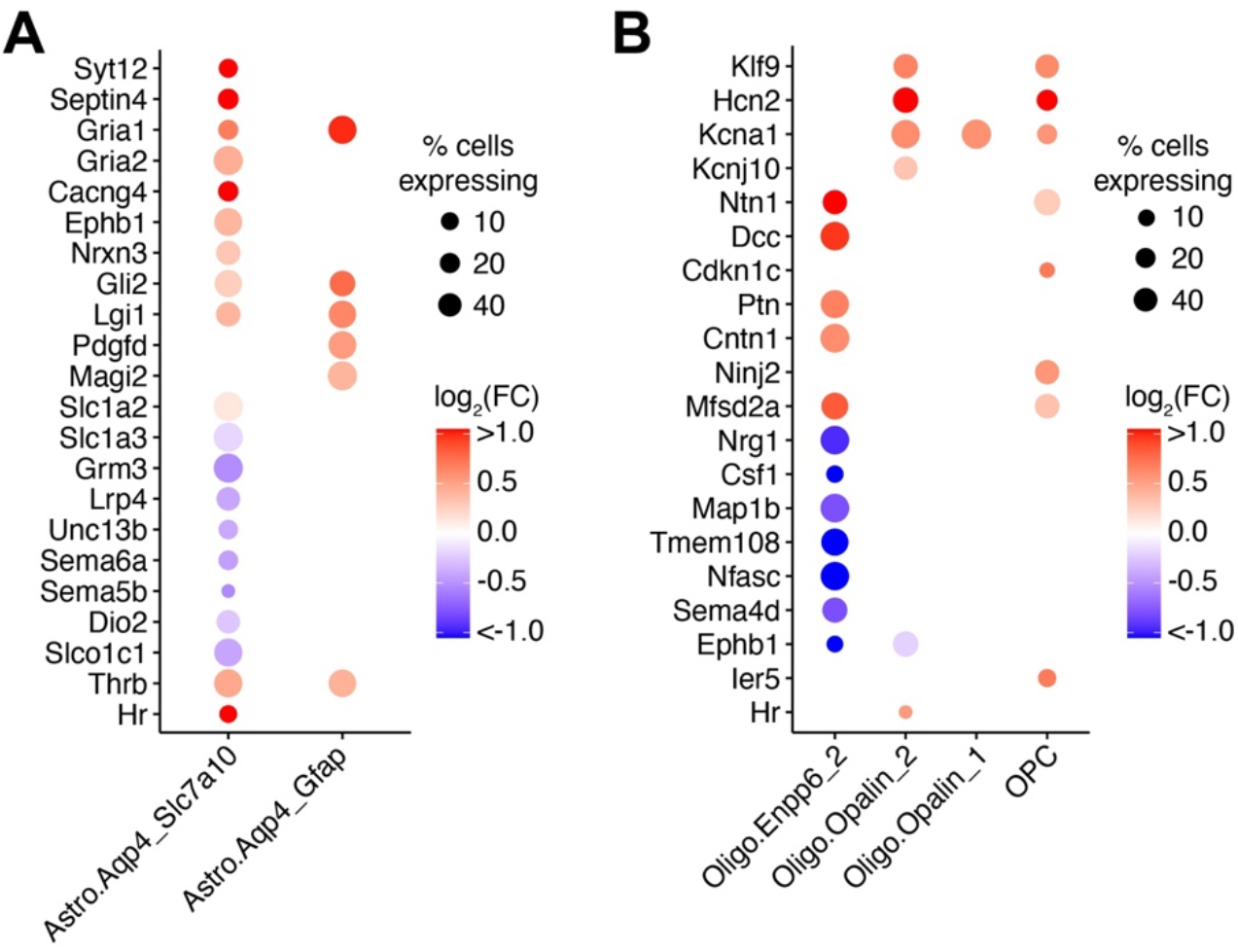
T3 activates genes associated with synapse regulation in astrocytes and differentiation and myelination in OPCs and oligodendrocytes. A) Dot plot highlighting TRGs that drive GSEA enriched pathways in astrocytes and are associated with astrocyte regulation and assembly of synapses. These include astrocyte expressed genes such as *Septin4*, *Slc1a2* and *Slc1a3*, which encode proteins implicated in glutamate clearance^116,117^, and *Lrp4*, which encodes an astrocytic modulator of glutamatergic synaptic release probability^118^. Additional TRGs are included that indicate a homeostatic response to T3 (downregulation of *Slco1a1* and *Dio2*, upregulation of *Thrb*). B) Dot plot highlighting TRGs that drive GSEA enriched pathways in OPCs and oligodendrocytes and are associated with OPC and oligodendrocyte differentiation, maturation, and myelination. These include genes encoding transcription factors such as *Klf9* known to drive oligodendrocyte differentiation and myelin regeneration^51^, and ion channels such as *Hcn2* that regulates myelin sheath length^119^, along with secreted factors such as *Ntn1*, which is implicated in oligodendrocyte maturation^120^ and is a ligand of the Robo3/DCC complex^54^. Dot color indicates the change in expression between the control and T3 conditions (log_2_(fold-change)). The size of the dot indicates the percentage of cells in each cell type that expressed one or more transcripts of the given gene in the T3 condition. Dots are only shown for genes in cell types where they were shown to be differentially expressed between control and T3 conditions with FDR-adjusted p < 0.05, and robustness score ≥ 0.5.

**Supplemental Figure 4.**
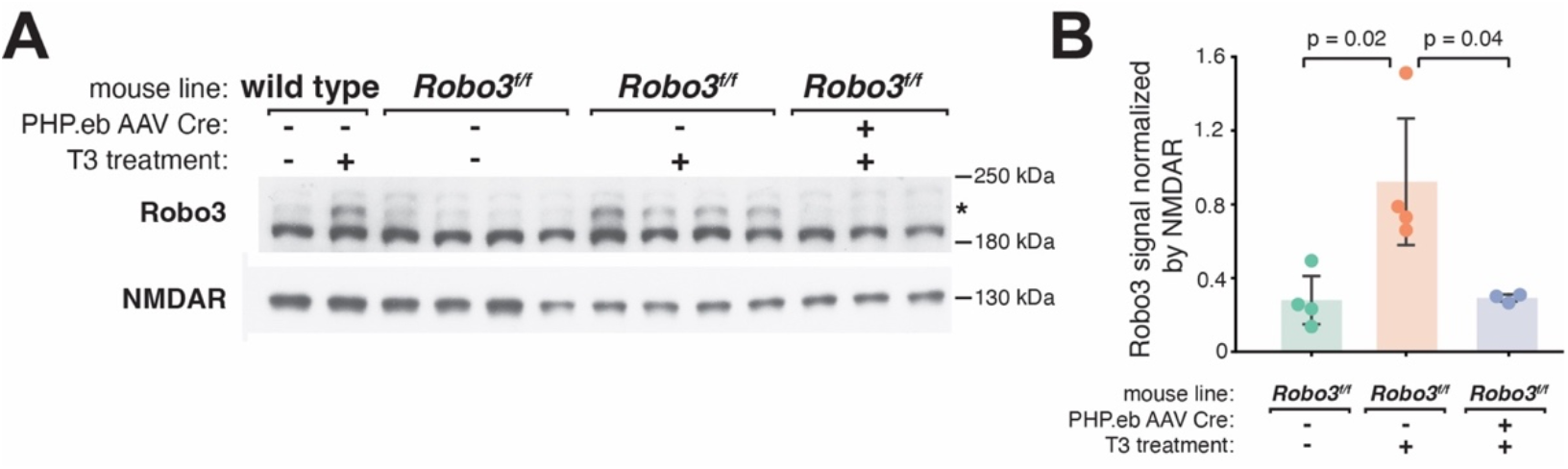
Robo3 protein levels increase with T3 in frontal cortex. A) Western blot from frontal cortex of wild type (left) and *Robo3^fl/fl^* animals with and without T3 treatment and/or Cre delivery (via systemic AAV delivery). Staining was performed with antibodies for Robo3, and the Glun1 subunit of the NMDA-type glutamate receptor to normalize for neuronal content in each sample. Robo3 antibody staining resulted in two bands, one of which increased in intensity with T3 treatment. We delivered Cre to excise Robo3 and prevent its induction with T3. Only the top band (*), whose intensity was altered by T3, was eliminated, demonstrating that this band is specific to Robo3, while the other band is non-specific. B) Robo3 protein levels increased with T3 treatment (p = 0.02) and were occluded by Cre excision of Robo3 (p = 0.04). Bars indicate mean, error bars are standard deviation, and p-values computed with two-tailed t-tests.

**Supplemental Figure 5.**
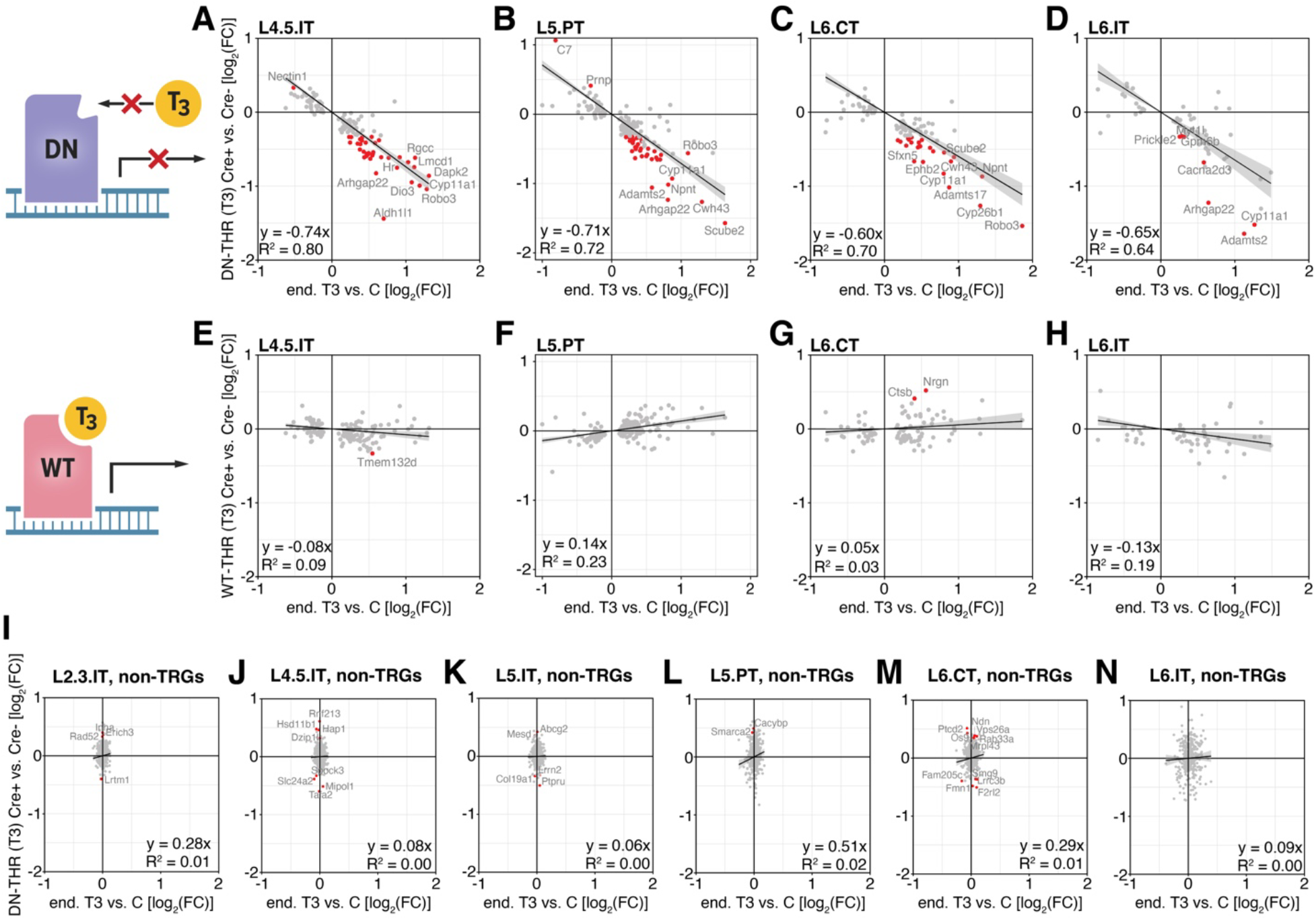
DN-THR selectively perturbs T3-induced genes, while WT-THR preserves TRG programs. A) Plots showing on the y-axis the log_2_(fold-change) of L4.5.IT TRGs (**Fig. 2b**) between Cre+ (DN-THR expressing) L4.5.IT cells and Cre-(lacking DN-THR) L4.5.IT cells, after T3 treatment. The x-axis shows the log_2_(fold-change) of L4.5.IT TRGs between the T3 and vehicle control conditions from the original dataset (**Fig. 2**). Red dots highlight TRGs whose expression was significantly disrupted by 25% or more due to DN-THR (FDR-adjusted p < 0.05, and fractional change in expression of at least ±25%). Linear regression fits to the data are overlaid, grey shading indicates a 95% confidence interval. Fit equation and R^2^ value are displayed on the lower left. B) As in **A)**, but for L5.PT neurons. C) As in **A)**, but for L6.CT neurons. D) As in **A)**, but for L6.IT neurons. E) As in **A)**, but for WT-THR expressing tissue. The y-axis shows the log_2_(fold-change) of L4.5.IT TRGs between Cre+ (WT-THR expressing) L4.5.IT cells and Cre-(lacking WT-THR) L4.5.IT cells, after T3 treatment. Red dots highlight TRGs as in **A)**. F) As in **E)**, but for L5.PT neurons. G) As in **E)**, but for L6.CT neurons. H) As in **E)**, but for L6.IT neurons. I) Plot showing on the y-axis the log_2_(fold-change) of L2.3.IT genes not regulated by T3 (non-TRGs) between Cre+ (DN-THR expressing) L2.3.IT cells and Cre-(lacking DN-THR) of L2.3.IT cells, after T3 treatment. The x-axis shows the log_2_(fold-change) of L2.3.IT non-TRGs (**Methods**) between the T3 and vehicle control conditions from the original dataset (**Fig. 2**). Red dots indicate non-TRGs whose expression was significantly disrupted by 25% or more due to DN-THR (FDR-adjusted p < 0.05, and fractional change in expression of at least ±25%). Linear regression fits to the data are overlaid, grey shading indicates a 95% confidence interval. Fit equation and R^2^ value are displayed on the lower right. J) As in **I)**, but for L4.5.IT neurons. K) As in **I)**, but for L5.IT neurons. L) As in **I)**, but for L5.PT neurons. M) As in **I)**, but for L6.CT neurons. N) As in **I)**, but for L6.IT neurons.

**Supplemental Figure 6.**
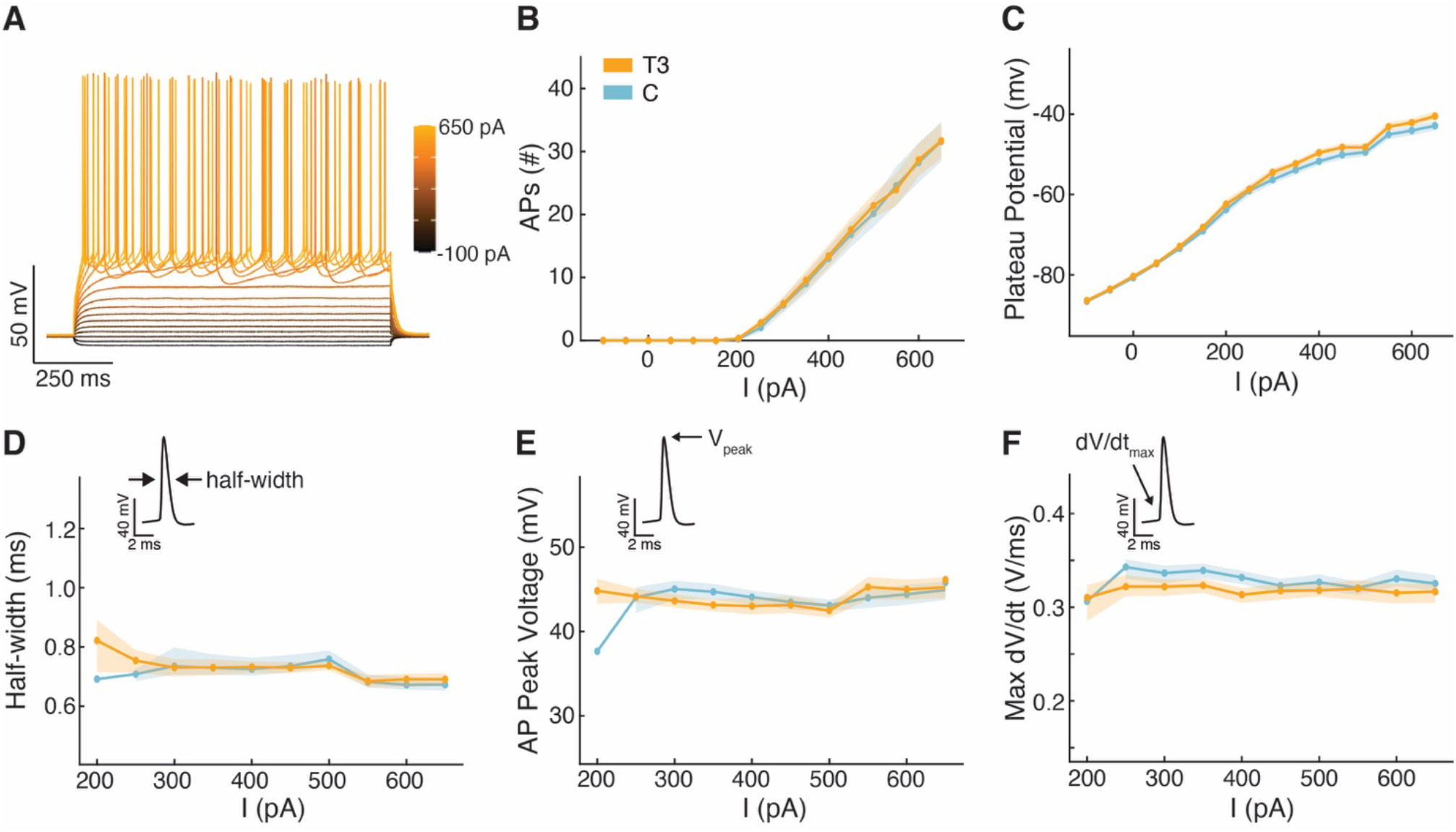
Intrinsic excitability of layer 2/3 pyramidal neurons is not modulated by T3. Whole-cell recordings were carried out on pyramidal neurons in M2 from T3 (n = 36 cells, 7 animals) or control (n = 30 cells, 7 animals) treated animals (3.5 days of treatment). Current steps (1 s) were injected from -100 pA to 650 pA. **A)** Example recording of a pyramidal neuron from a T3-treated animal. Traces are color coded by the injected current. **B)** Input-output curve of injected current vs. number of generated action potentials (APs). There was no observed effect on the generation of APs due to T3-treatment (p = 0.85). **C)** Plateau potential (steady-state elevated baseline potential) for each current step. There was no observed difference in plateau potentials due to treatment (p = 0.17). **D)** Half-width (width of the action potential at 50% of its peak magnitude) of the first generated AP for each current step. Inset: schematic of the half-width measurement. There was no observed difference in half-width due to treatment (p = 0.73). **E)** Peak voltage of the first generated AP for each current step. There was no observed effect of T3-treatment (p = 0.85). **F)** Maximum rate of change in voltage (dV/dt) of the first generated AP for each current step. There was no observed effect of T3-treatment (p = 0.43). Experiments were repeated using alternate divalent cation concentrations (1 mM Ca^2+^, 2 mM Mg^2+^). Comparison between control (n = 10 cells, 2 animals) and T3-treated (n = 11 cells, 2 animals) did not reveal any modulation by T3 (I vs. AP: p = 0.99; I vs. plateau: p = 0.76; I vs. half-width: p = 0.43; I vs. peak: p = 0.70; I vs. max dV/dt: p = 0.55). All p-values were calculated by likelihood ratio tests comparing linear mixed-effects models (LMMs) including the treatment condition vs. LMMs that did not.

**Supplemental Figure 7.**
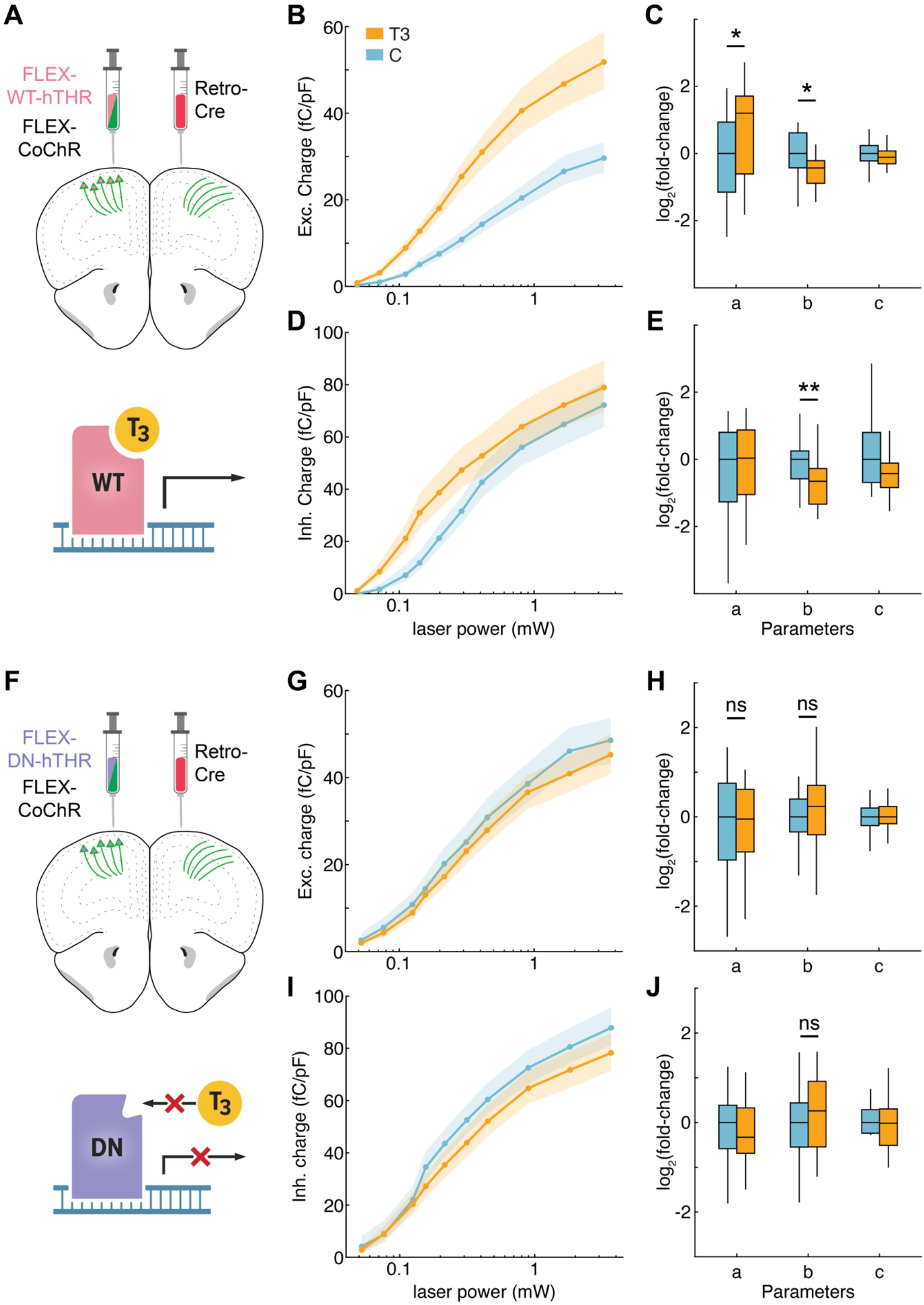
Presynaptic actions of T3. A) Top: Injection schematic. AAVs encoding a Cre-dependent CoChR (FLEX-CoChR) and a Cre-dependent WT-THR (FLEX-WT-hTHR) were co-delivered to the upper layers of M2. A retrograde AAV encoding Cre was delivered to the contralateral hemisphere, resulting in CoChR and nuclear WT-THR expression selectively in neurons sending projections to contralateral M2. Bottom: Cartoon of THR expression. B) Normalized post-synaptic excitatory charge as a function of laser stimulus power from experiments including WT-THR expression in the presynaptic neurons. Orange are measures from T3-treated mice (n = 27 neurons, 10 mice), and blue are measures from vehicle treated mice (n = 30 neurons, 9 mice). Dots represent mean values; shading indicates bootstrapped standard error of the mean. C) Boxplot of changes in each sigmoid parameter (from single cell fits of excitatory charges vs. laser power curves) relative to the median control value, for experiments including WT-THR expression in the presynaptic neurons. The saturation amplitude (a) was significantly increased by T3-treatment (p = 0.026), and the power to half-maximum (b) was significantly decreased by T3-treatment (p = 0.014). D) As in B), but for light-evoked disynaptic inhibitory post-synaptic currents (IPSCs). E) As in C), but for single cell fits of inhibitory charge vs. laser power curves. The power to half-maximum was significantly decreased by T3-treatment (p = 0.004). F) Top: As in A), but for experiments that replaced FLEX-WT-hTHR with FLEX-DN-hTHR, an AAV encoding a Cre-dependent DN-THR. Bottom: Cartoon of DN-THR expression which perturbs T3-dependent gene transcription. G) As in B), but for experiments with DN-THR expressing in presynaptic neurons. Orange are measures from T3-treated mice (n = 31 neurons, 8 mice), and blue are measures from vehicle treated mice (n = 31 neurons, 8 mice). H) As in C), but for experiments with DN-THR expressing in presynaptic neurons. The saturation amplitude (a) was not altered by T3-treatment (p = 0.96), and the power to half-maximum (b) was not altered by T3-treatment (p = 0.521). I) As in G), but for light-evoked inhibitory post-synaptic currents (IPSCs). J) As in H), but for single cell fits of inhibitory charge vs. laser power curves. The power to half-maximum was not altered by T3-treatment (p = 0.36). All statistical comparisons performed with the Mann Whitney U test. * p < 0.05, ** p < 0.01.

**Supplemental Figure 8.**
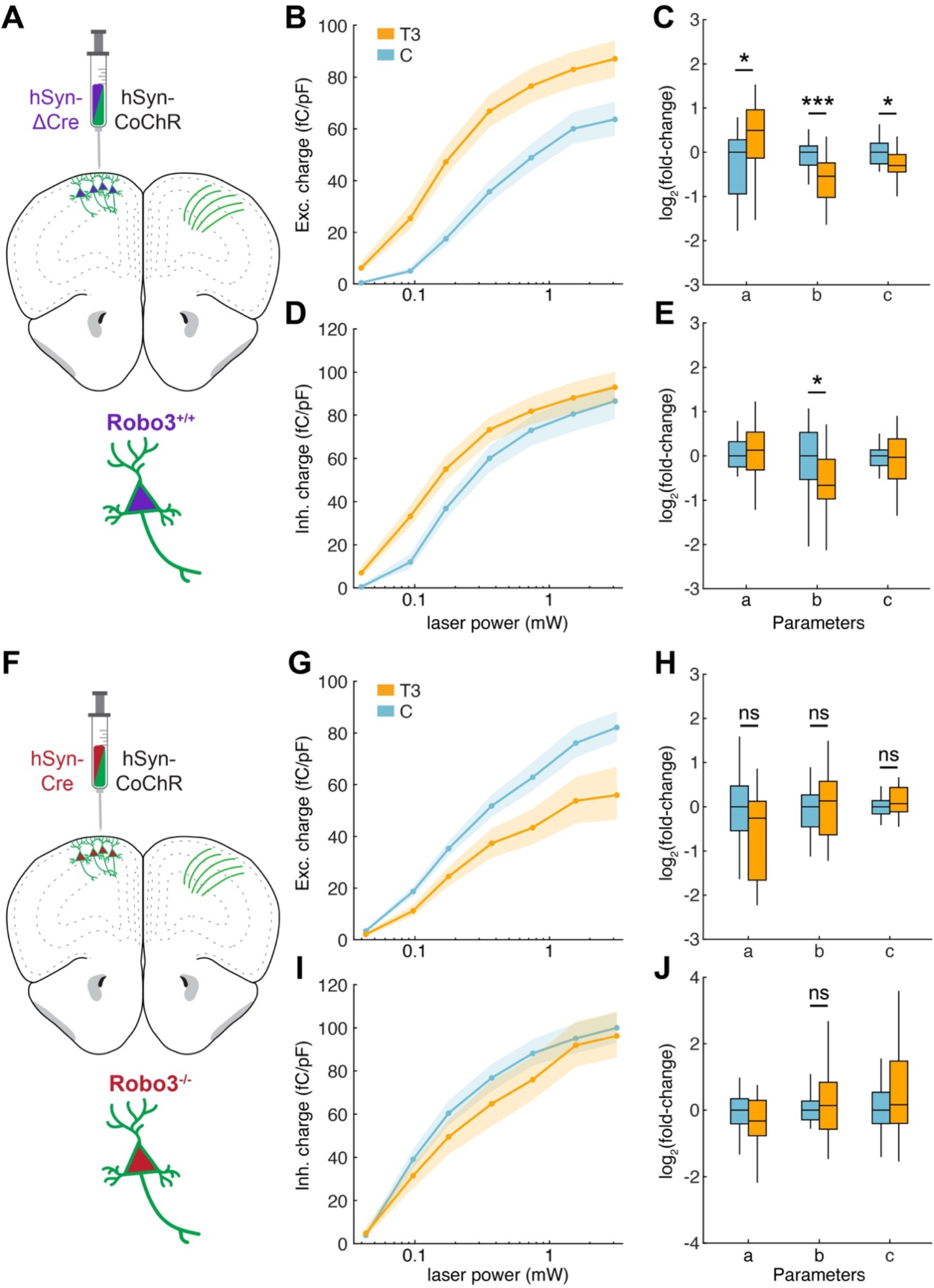
Robo3 is required to observe T3-dependent changes in synaptic connectivity. A) Top: Injection schematic. AAVs encoding an inactive control ΔCre (incapable of catalyzing recombination, hSyn-ΔCre) and a Cre-independent CoChR-GFP (hSyn-CoChR), both driven by neuronal specific promoter hSyn, were co-delivered to the upper layers of M2. Whole cell electrophysiology recordings were made from the contralateral M2 by stimulating labeled axons containing CoChR-GFP and inactive ΔCre from cortico-cortical projecting IT neurons. Bottom: Cartoon of a cortico-cortical projecting IT neuron expressing ΔCre, leaving *Robo3* expression intact. B) Normalized post-synaptic excitatory charge as a function of laser stimulus power from experiments with presynaptic ΔCre and CoChR-GFP expression. Orange are measures from T3-treated mice (n = 33 neurons, 7 mice), and blue are measures from vehicle treated mice (n = 17 neurons, 4 mice). Dots represent mean values; shading indicates bootstrapped standard error of the mean. C) Boxplot of changes in each sigmoid parameter relative to the median control value, for experiments including ΔCre expression in the presynaptic neurons. These experiments recapitulated previously observed changes due to T3. The saturation amplitude (a) was significantly increased by T3 treatment (p = 0.017), and the power to half-maximum (b) was significantly decreased by T3 treatment (p < 0.001). The slope of the sigmoid was also significantly decreased by T3 treatment (p = 0.01) consistent with the trend observed in previous experiments (**Fig. 4e, Supp. Fig. 7c**) D) As in **B)**, but for light-evoked inhibitory post-synaptic currents (IPSCs). E) As in **C)**, but for single cell fits of inhibitory charge vs. laser power curves. Experiments with βCre expression in the presynaptic neurons recapitulated previous findings of T3-dependent changes to IPSCs. The power to half-maximum was significantly decreased by T3-treatment (p = 0.013). A) F) Top: As in **A)**, but for experiments that replaced the inactive control ΔCre with a functional Cre (hSyn-Cre), leading to the loss of Robo3 expression (**Supp. Fig. 3**) Bottom: Cartoon of a cortico-cortical projecting IT neuron expressing Cre, leading to the loss of *Robo3* expression. B) G) As in **B)**, but for experiments with Cre expressing in presynaptic neurons. Orange are measures from T3-treated mice (n = 17 neurons, 5 mice), and blue are measures from vehicle treated mice (n = 33 neurons, 7 mice). C) H) As in **C)**, but for experiments with Cre expressing in presynaptic neurons. The saturation amplitude (a) was no longer altered by T3-treatment (p = 0.06), the power to half-maximum (b) was not altered by T3-treatment (p = 0.95), and the slope was not altered by T3-treatment (p = 0.57). I) As in **G)**, but for light-evoked inhibitory post-synaptic currents (IPSCs). D) J) As in **H)**, but for single cell fits of inhibitory charge vs. laser power curves. The power to half-maximum was no longer altered by T3-treatment (p = 0.76). All statistical comparisons performed with the Mann Whitney U test. * p < 0.05, ** p < 0.01.

**Supplemental Figure 9.**
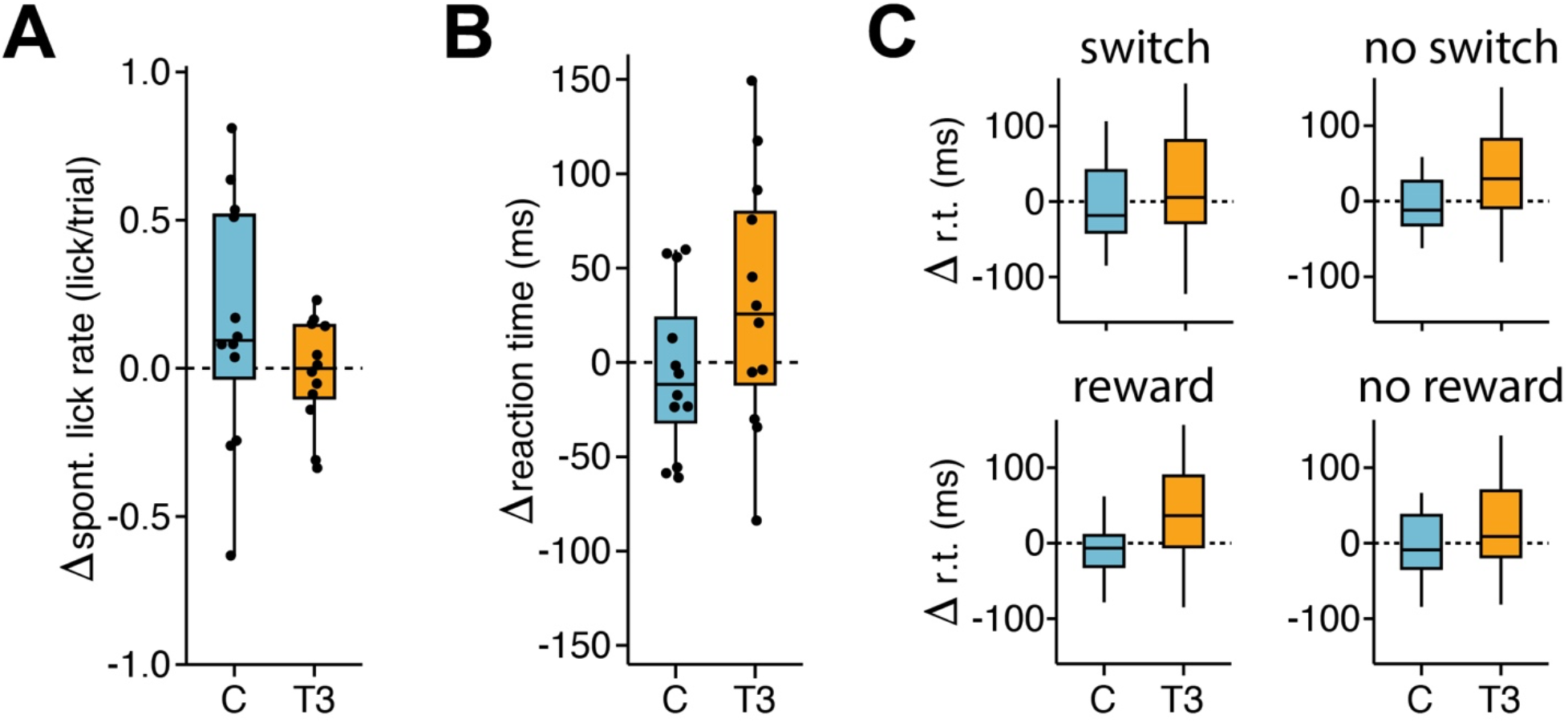
Gross motor actions are not altered by T3. A) Change in the spontaneous lick rate (licks/trial) between the habituation period and days 4-7 of treatment for each experimental cohort (control cohort, blue; T3 cohort, orange). The spontaneous lick rate is defined as the average number of licks during the un-cued no-lick period that precedes the auditory tone for each trial. To exclude lick bouts due to consumption of a reward, only trials in which the previous trial was unrewarded are included in this analysis. Neither cohort had a significant change in spontaneous lick rate (control cohort, p = 0.23; T3 cohort, p = 0.77, t-test). B) Change in the reaction time (ms) of the selection lick after the auditory tone (measure as the time between tone onset and lick contact with the spout) between the habituation period and days 4-7 of treatment for each experimental cohort. Neither cohort had a significant change in reaction time (control cohort, p = 0.70; T3 cohort, p = 0.14, t-test). C) Change in the reaction time (ms) between the habituation period and days 4-7 of treatment for each experimental cohort conditional on whether the trial resulted in a switch in motor action (top row), or whether the column followed a rewarded or unrewarded trial (bottom row). Neither cohort had a significant change in any of the conditional reaction times (switch: p = 0.96 for C, p = 0.64 for T3; no switch: p = 0.71 for C, p = 0.12 for T3; previous trial rewarded: p = 0.63 for C, p = 0.12 for T3; previous trial unrewarded: p = 0.77 for C, p = 0.17 for T3; t-test). Black dots represent single mice. For all analyses, n = 12 animals for each treatment condition (T3 or control).

**Supplemental Figure 10.**
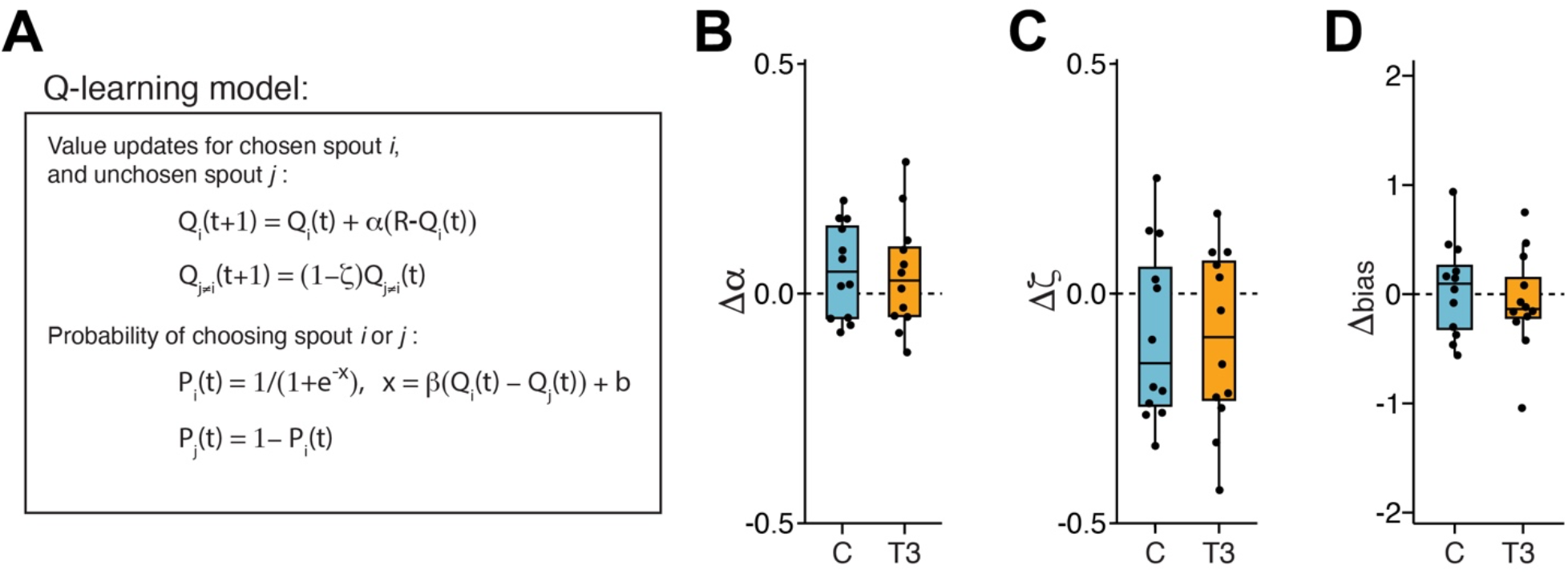
Q-learning model for 2ABT task. A) Update equations governing action values for each spout (top pair of equations). For the chosen spout *i* (either left or right), the value of selecting the spout on the next trial, Q_i_(t+1), is updated by the difference between the reward (R = 1 for a dispensed water droplet, R = 0 for a failure) and current value of the spout, Q_i_(t), scaled by the learning rate α. The value of the unchosen spout (*j* ≠ *i*, i.e. left spout if right spout was chosen) for the next trial, Q_j_(t+1), is a decremented value of the current value, Q_j_(t), scaled by the forgetting rate ζ. The Q-values are then fed into a Boltzmann distribution (softmax, bottom pair equations) which determines the policy (probability) of selecting each spout. β is an inverse temperature parameter that determines the degree of exploration vs. exploitation given relative values between the spouts Q_i_(t)-Q_j_(t). b is a fixed bias parameter. The model fit the data well for all treatments and epochs (spout-choice prediction accuracy on held-out data during habituation for T3 cohort: 0.85 ± 0.03, mean ± std. dev.; and during days 4-7 of treatment for T3 cohort: 0.85 ± 0.03; comparison between epochs: p = 0.52, t-test. Spout-choice prediction accuracy on held-out data during habituation for C cohort: 0.85 ± 0.03; and during days 4-7 of treatment for C cohort: 0.86 ± 0.03; comparison between epochs: p = 0.64, t-test). A) B) Change in the learning rate parameter α between the habituation period and days 4-7 of treatment for each experimental cohort (control cohort, blue; T3 cohort, orange). Black dots represent single mice. Neither cohort had a significant change in α (control cohort, p = 0.11; T3 cohort, p = 0.28, t-test). B) Change in the forgetting rate parameter ζ between the habituation period and days 4-7 of treatment for each experimental cohort (control cohort, blue; T3 cohort, orange). Black dots represent single mice. Neither cohort had a significant change in α (control cohort, p = 0.14; T3 cohort, p = 0.10, t-test). C) Change in the bias parameter b between the habituation period and days 4-7 of treatment for each experimental cohort (control cohort, blue; T3 cohort, orange). Black dots represent single mice. Neither cohort had a significant change in α (control cohort, p = 0.70; T3 cohort, p = 0.64, t-test). For all analyses, n = 12 animals for each treatment condition (T3 or control). Analysis of the changes in the β parameter appear in **Fig. 5j**.

**Supplemental Figure 11.**
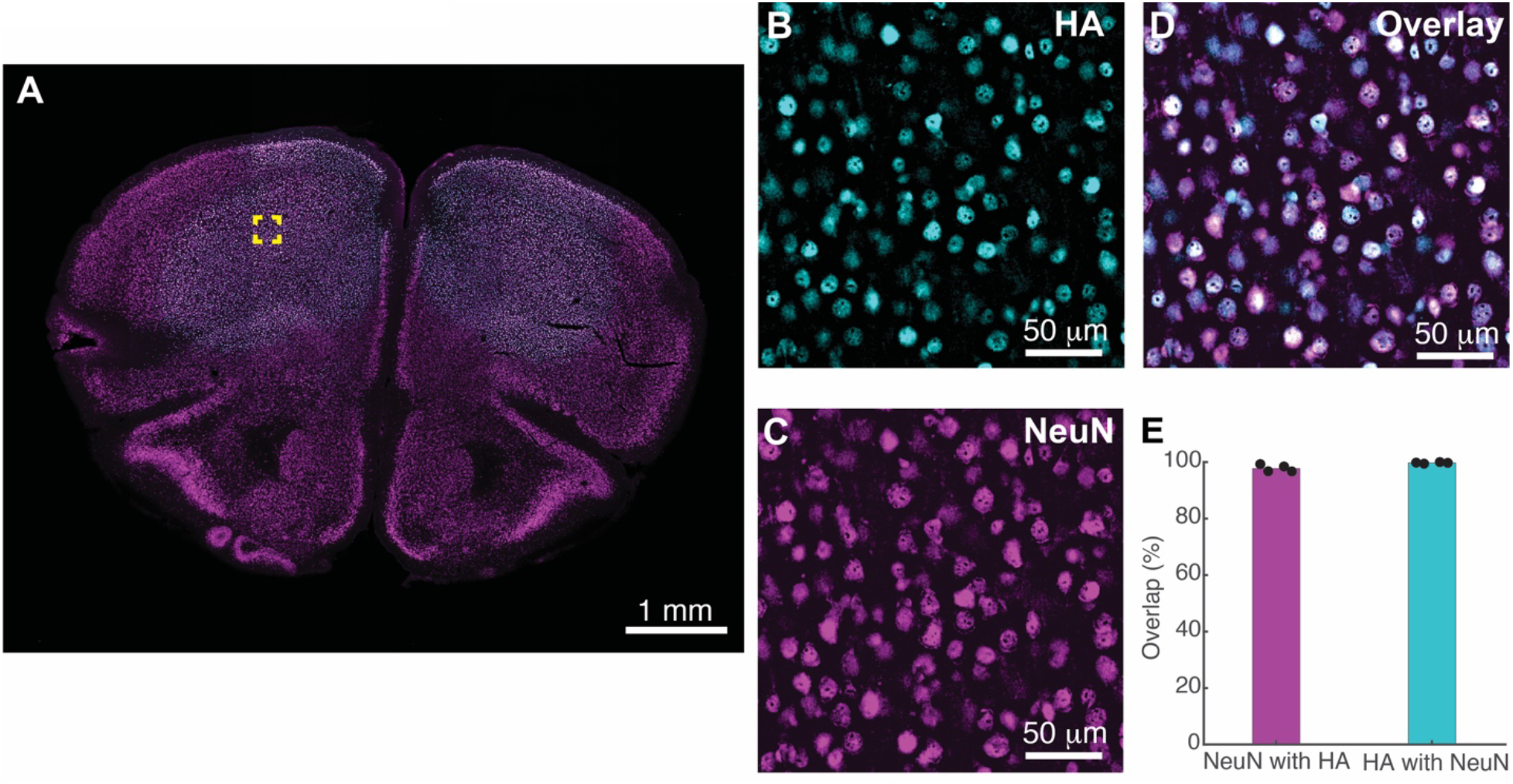
AAV mediated expression of hSyn-THR is specific to neurons. A) Low magnification image showing expression of WT-THR from AAV injections of hSynapsin-1 driven transgene expression (Cre-independent) in frontal cortex (∼2.5 mm anterior to bregma). Immunohistochemistry was performed with antibodies for HA (WT- and DN-THR constructs have a C-terminal HA tag, cyan) and neuronal peri-nuclei (NeuN, magenta). B) High magnification image of HA staining from the yellow region highlighted in A). C) As in **B**) with NeuN staining. D) Overlay of images from **B**) and **C**). E) Quantification of HA and NeuN overlap. HA was present in 97.78% (1274/1302 cells) of NeuN+ peri-nuclei, and 99.72% (1522/1527 cells) of HA+ nuclei had NeuN expression. Bar graphs represent the ratio of HA+ / NeuN+ cells and NeuN+ / HA+ cells. Bars indicate mean, black dots indicate individual samples (2 hemispheres from 2 animals, 4 samples total). Each quantification was validated near the center of the injection site.

**Supplemental Figure 12.**
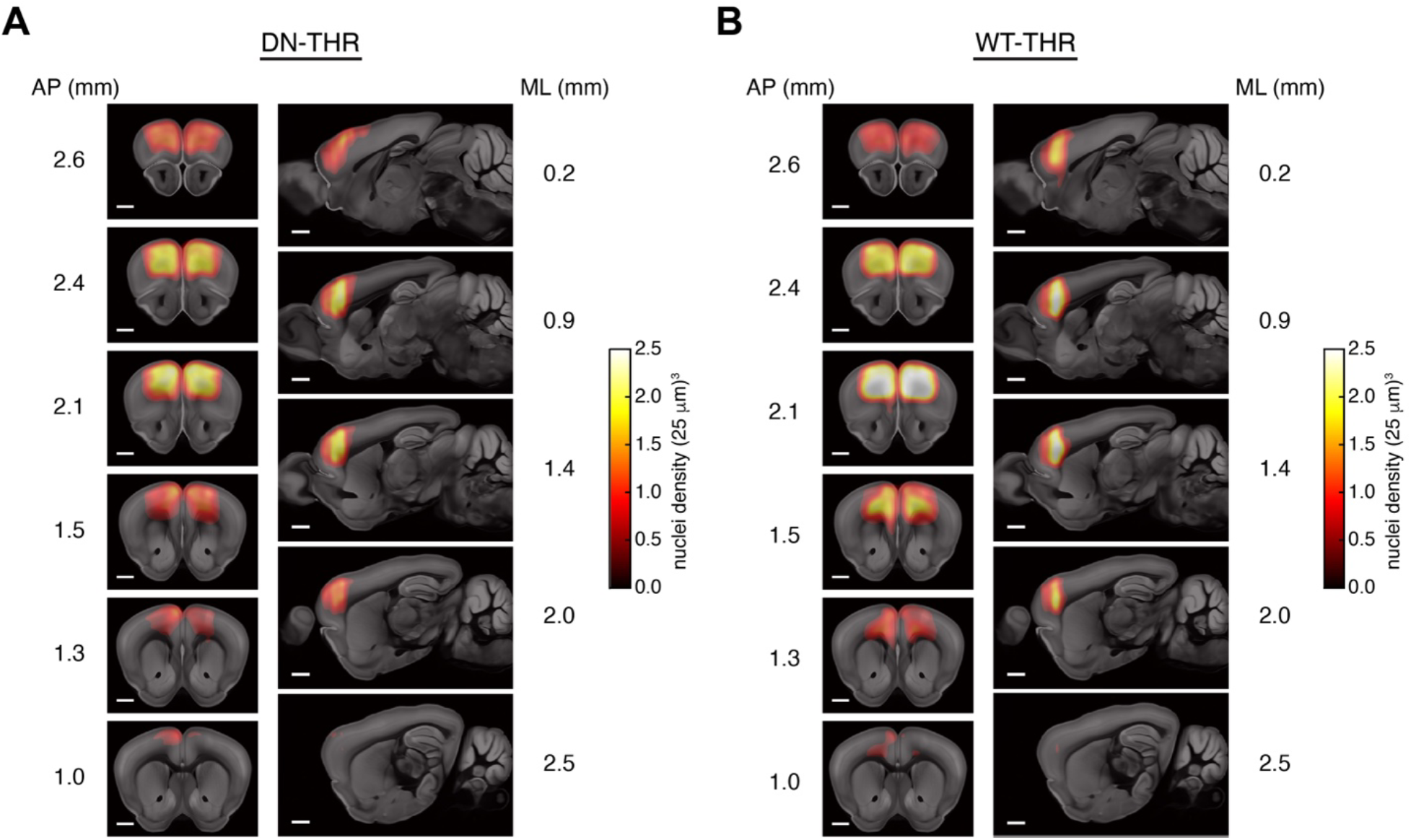
WT-/DN-THR expression heatmaps. A) Coronal (left) and sagittal (right) sections characterizing average DN-THR expression across the cohort (n = 10 animals). A full list of counts per brain region are in **Supp. Table 4**. B) As in panel (A) for the WT-THR cohort (n = 12 animals). A full list of counts per brain region are in **Supp. Table 4**. AP = anterior/posterior axis, measures relative to bregma. ML = medial/lateral axis, measures relative to the midline. All scale bars are 1 mm.

**Supplemental Figure 13.**
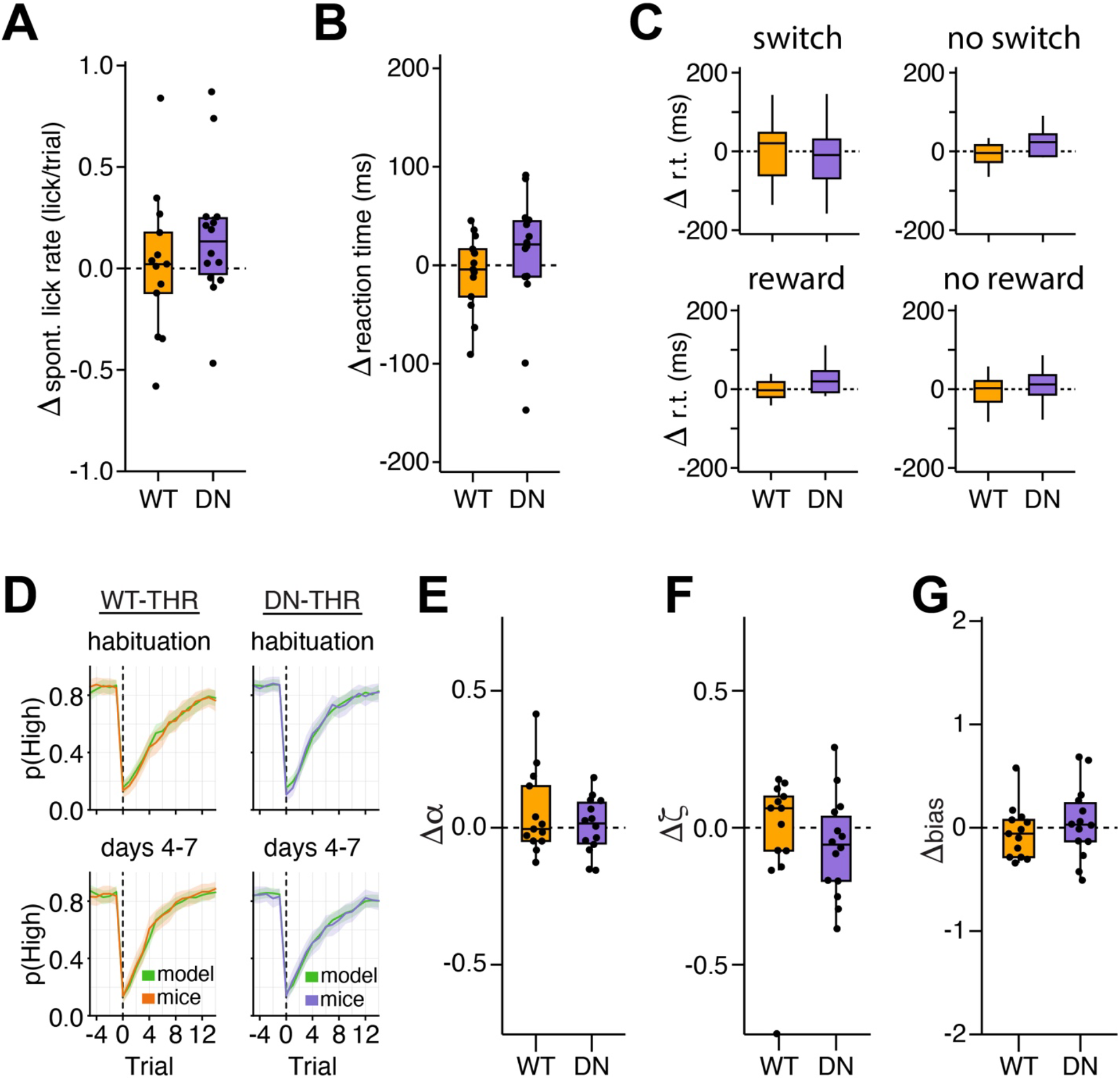
WT-/DN-THR motor action measures and Q-learning parameters. A) Change in the spontaneous lick rate (as in **Supp. Fig. 9**) between the habituation period and days 4-7 of treatment for each experimental cohort (WT-THR cohort, orange; DN-THR cohort, purple). Neither cohort had a significant change in spontaneous lick rate (WT-THR cohort, p = 0.82; DN-THR cohort, p = 0.10, t-test). B) Change in the reaction time (ms) of the selection lick after the auditory tone between the habituation period and days 4-7 of treatment for each experimental cohort. Neither cohort had a significant change in reaction time (WT-THR cohort, p = 0.46; DN-THR cohort, p =0.65, t-test). C) Change in the reaction time (ms) between the habituation period and days 4-7 of treatment for each experimental cohort conditional on whether the trial resulted in a switch in motor action (top row), or whether the column followed a rewarded or unrewarded trial (bottom row). Neither cohort had a significant change in any of the conditional reaction times (switch: p = 0.88 for WT-THR, p = 0.66 for DN-THR; no switch: p = 0.34 for WT-THR, p = 0.27 for DN-THR; previous trial rewarded: p = 0.44 for WT-THR, p = 0.47 for DN-THR; previous trial unrewarded: p = 0.77 for WT-THR, p = 0.58 for DN-THR; t-tests or Wilcox signed rank tests dependent on whether distributions were significantly non-gaussian). D) Q-learning model fits of the probability of selecting the highly rewarding spout, p(High). Data from the habituation period (top row) and days 4-7 (bottom row). Orange/purple lines are the mean probability from the mouse data (WT-/DN-THR respectively), and the green line is the model prediction. Shaded areas indicate 95% confidence intervals. The model fit the data well for all treatments and epochs (spout-choice prediction accuracy on held-out data during habituation for WT-THR cohort: 0.88 ± 0.03, mean ± std. dev.; and during days 4-7 of treatment for WT-THR cohort: 0.87 ± 0.03; comparison between epochs: p = 0.21, t-test. Spout-choice prediction accuracy on held-out data during habituation for DN-THR cohort: 0.87 ± 0.04; and during days 4-7 of treatment for DN-THR cohort: 0.84 ± 0.05; comparison between epochs: p = 0.13, t-test). 1. E) Change in the learning rate parameter α between the habituation period and days 4-7 of treatment for each experimental cohort. Neither cohort had a significant change in α (WT-THR cohort, p = 0.23; DN-THR cohort, p = 0.79, t-test). 2. F) Change in the forgetting rate parameter ζ between the habituation period and days 4-7 of treatment for each experimental cohort. Neither cohort had a significant change in ζ (WT-THR cohort, p = 0.68; DN-THR cohort, p = 0.18, t-test). 3. G) Change in the bias parameter ‘b’ between the habituation period and days 4-7 of treatment for each experimental cohort. Neither cohort had a significant change in ‘b’ (WT-THR cohort, p = 0.52; DN-THR cohort, p = 0.59, t-test). Black dots represent single mice. For all analyses, WT-THR cohort: n = 13 animals, DN-THR cohort: n = 14 animals. Analysis of the changes in the β parameter appear in **Fig. 6i**.

**Supplemental Table 1.** TRGs that were differentially expressed between control and elevated T3 conditions (FDR-adjusted p < 0.05, robustness score ≥ 0.5) for each cell type.

**Supplemental Table 2.** Gene set enrichment analyses for each cell type.

**Supplemental Table 3.** Cell properties of L2/3 pyramidal neurons across conditions and experiments (Fig. 4, Supp. Fig 7-8). Values are mean +-sem. Statistical comparisons performed with Mann Whitney U test.

| Experiment | C<sub>membrane</sub><br>(pF) | C<sub>membrane</sub><br>(p-value) | R<sub>membrane</sub><br>(MΩ) | R<sub>membrane</sub><br>(p-value) | R<sub>series</sub><br>(MΩ) | R<sub>series</sub><br>(p-value) | I<sub>hold</sub> (pA) | I<sub>hold</sub><br>(p-value) |
| --- | --- | --- | --- | --- | --- | --- | --- | --- |
| No<br>exogenous<br>expression<br>(Fig 4) | C: 218 ±<br>65<br>T3: 197<br>± 55 | p = 0.26 | C: 131 ±<br>57<br>T3: 130<br>± 49 | p = 0.98 | C: 10.3 ±<br>3.9<br>T3: 10.4<br>± 3.5 | p =<br>0.65 | C: -118<br>± 89<br>T3: -106<br>± 82 | p =<br>0.67 |
| WT-THRB<br>(Supp. Fig 7) | C: 227 ±<br>52<br>T3: 214<br>± 47 | p = 0.33 | C: 124 ±<br>51<br>T3: 121<br>± 43 | p = 0.93 | C: 8.9 ±<br>2.6<br>T3: 9.1 ±<br>2.9 | p =<br>0.73 | C: -75 ±<br>70<br>T3: -121<br>± 94 | p =<br>0.054 |
| DN-THRB<br>(Supp. Fig 7) | C: 233 ±<br>56<br>T3: 241<br>± 60 | p = 0.61 | C: 115 ±<br>39<br>T3: 118<br>± 42 | p = 0.89 | C: 7.8 ±<br>2.0<br>T3: 8.3 ±<br>2.7 | p =<br>0.29 | C: -104<br>± 103<br>T3: -120<br>± 97 | p =<br>0.31 |
| ΔCre,<br>Robo3<sup>+/+</sup><br>(Supp. Fig 8) | C: 228 ±<br>43<br>T3: 227<br>± 52 | p = 0.87 | C: 114 ±<br>29<br>T3: 102<br>± 42 | p = 0.09 | C: 10.2 ±<br>3.2<br>T3: 9.9 ±<br>3.5 | p =<br>0.61 | C: -130<br>± 93<br>T3: -101<br>± 92 | p =<br>0.21 |
| Cre,<br>Robo3<sup>-/-</sup><br>(Supp. Fig 8) | C: 264 ±<br>77<br>T3: 240<br>± 66 | p = 0.31 | C: 94 ±<br>41<br>T3: 99 ±<br>36 | p = 0.46 | C: 8.1 ±<br>2<br>T3: 9.8 ±<br>3.2 | p =<br>0.06 | C: -97 ±<br>87<br>T3: -102<br>± 119 | p =<br>0.68 |

**Supplemental Table 4.** Mean counts (and standard error of the mean) of detected nuclei across brain regions expressing DN-THR (n = 10 animals) or WT-THR (n = 12 animals). Also includes normalized counts (normalized to root for each brain).

## Author contributions

Conceptualization: D.R.H., B.L.S.

Performed research: D.R.H., A.C.D., H.C.F., L.H., G.K., W.W., K.R., C.P., A.S., Y.Y., C.B, N.M.N, S.P., G.L.B., A.G., M.C., L.C.

Analysis: D.R.H, A.U., G.K., A.S.

Supervision: D.R.H., B.L.S, E.Z.M., G.L.B., M.E.G., A.B.

Reagents: T.S.

Contributed software: D.R.H., R.H., C.C.B, A.E.G., L.G. Writing: D.R.H., B.L.S.

Funding: D.R.H, B.L.S.

All authors gave feedback on the manuscript.

## Acknowledgments

We thank Anthony Hollenberg, Kristen Vella, Minsuk Hyun, Kee Wui Huang, and members of the Sabatini lab for helpful conversations, advice, and technical assistance. We thank Alain Chédotal for *Robo3^fl/fl^* mice. We thank Barbara Caldarone and the Harvard Medical School Mouse Behavior Core for assistance and facility use for LD experiments. We thank the Harvard Medical School Research Instrumentation Core, Paul Jonak, and Gil Mandelbaum for assistance in designing and implementing the 2ABT hardware and software. We thank Nina Sachdev for assistance with the single cell data analysis. This work was supported by a Burroughs Wellcome Fund Career at the Scientific Interface Award, a Baszucki Brain Research Fund Bipolar Disorder Grant, a Blavatnik Biomedical Accelerator Grant, and NIH (R37NS046579). A.U. is supported by a T32GM144273 from the National Institute of General Medical Sciences and T32HG002295 from the National Human Genome Research Institute, NIH.

## Methods

### Mice

The following mice were used: C57BL6/J (Jackson labs #000664), and *Robo3^fl/fl^* (gift from Alain Chédotal), which were bred on a C57BL6/J genetic background. Sex-dependent effects of thyroid hormone have been reported across many animal species^17,121^, including mice^122^ and humans^102^. We therefore restricted ourselves to only males for these initial studies. All animal care and experimental manipulations were performed in accordance with protocols approved by the Harvard Standing Committee on Animal Care, following guidelines described in the US NIH Guide for the Care and Use of Laboratory Animals

### T3, sobetirome, and PTU delivery

Stock T3 (Sigma, T6397) solution was dissolved at 10 mg/mL in 100 mM NaOH and stored at -80° C. T3 working solution was prepared by making a 200x dilution of the stock in a 0.5% tween-20 solution in PBS for a final concentration of 50 μg/mL. A matching volume of 100 mM HCL was added to balance pH. The control solution was prepared identically substituting stock T3 with 100 mM NaOH, diluted 200x into the vehicle (0.5% tween-20 solution in PBS). Animals were weighed and appropriate volumes were delivered by twice daily intraperitoneal (IP) injection to achieve the desired concentration. A concentration of 0.125 μg/g was used unless otherwise noted.

The vehicle solution for sobetirome^45^ (from Thomas Scanlan) experiments was made by combining Kolliphor (sigma, C5135), NMP (Sigma, 328634), and water in a 1:1:8 ratio (KNH). Stock sobetirome solution was made by dissolving sobetirome in KNH at a final concentration of 3 mg/mL and stored at -80 °C. Sobetirome was diluted 100x into KNH for working solution and delivered by twice daily subcutaneous injection at 1 μg/g unless otherwise noted. The control solution was the vehicle KNH.

Mice were given 2 days of vehicle control injections for habituation. Mice were then injected with either the experimental solution or control solution twice per day. Experiments for the analysis of RNA were performed on the third day of treatment, after morning injection, unless otherwise noted. Experiments for the analysis of protein, electrophysiology, and behavior (light-dark paradigm) were performed on the fourth day of treatment, after morning injection, unless otherwise noted. For indirect calorimetry experiments of T3, treatment continued for 5 or more days (**Fig. 1c**). For the 2ABT (**Fig. 5-6**), treatment continued for 8 days total (days 0-7 of treatment).

For PTU treatment, mice were transitioned to an iodine deficient diet containing PTU (0.15% PTU; Inotiv, TD.95125) or a control diet (Inotiv, TD.97350) for 3 weeks. Although PTU mice received no injection, mice were handled twice a day for five days to provide similar habituation. On the sixth day of handling, mice were tested in the light-dark paradigm, and afterwards tissue was collected for RNA analysis.

### Quantitative PCR

Animals were anesthetized by isoflurane inhalation and trans-cardially perfused with an ice cold choline solution containing (in mM) 25 sodium bicarbonate, 12 glucose, 1.25 sodium phosphate monobasic monohydrate, 7.5 magnesium chloride hexahydrate, 2.5 potassium chloride, 10 HEPES, 110 choline chloride, 11.6 sodium L-ascorbate, 3 sodium pyruvate, pH 7.4. Dorsal anterior cortex and liver tissues were dissected from animals and stored at -80 °C. Samples were suspended and frozen in Trizol (Life Technologies, 15596018) before RNA was extracted following the RNEasy Micro Kit (Qiagen, 74004) protocol. RNA concentrations were measured by a Nanodrop spectrophotometer (ThermoFisher, ND-2000), and samples were diluted to a 50 ng/µl RNA concentration. cDNA was generated from 1 µl of diluted RNA sample using SuperScript IV VILO Master Mix Kit with ezDNase enzyme (ThermoFisher, 11766050). Quantitative PCR was performed with TaqMan probes for target genes *Hr* (ThermoFisher, Mm00498963_m1), *Dio1* (ThermoFisher, Mm00839358_m1), and *Ier5* (ThermoFisher, Mm01295615_s1) using a standard protocol on a QuantStudio 3 (ThermoFisher, A28567). *Gapdh* (ThermoFisher, Mm99999915_g1) served as a normalization factor for all gene probes. A standard curve using ten-fold dilutions was generated for both the gene-of-interest and *Gapdh* samples. Gene-of-interest C_t_ values were normalized to their respective *Gapdh* counterparts.

### FISH

Fresh-frozen mouse brain tissue was cryosectioned on a LEICA CM3050 S cryostat into 15µm sections, mounted on glass slides, allowed to dry at -20 °C for >20 minutes, and stored at -80 °C. 3- or 4-plex Fluorescent RNA in situ hybridization (FISH) was performed using Advanced Cell Diagnostics (ACD) RNAscope Multiplex Fluorescent Reagent Kit v1 (discontinued) or v2 (#323270 and #323120). Sections were prepared, pretreated, and processed according to ACD protocol except for the protease treatment, in which sections were treated with ACD protease III (#322337) for 10 minutes at room temperature. Target probes included *Hr* (#883311), *Ier5* (#530401-C3), and *Robo3* (#558811-C2). The fluorophores used were Opal 520 reagent (Akoya Biosciences, #OP-001001), Opal 570 reagent (#OP-001003), Opal 620 reagent (#OP-001004), and Opal 690 reagent (#OP-001006). Nuclei were stained using the ACD supplied DAPI stain for 30 s at RT and the slides were mounted using Fluromount-G (SouthernBiotech, # 0100-01). ACD’s 4-plex positive (#321811) and negative control (#321831) probes were used for experimental signal verification.

Samples were imaged on a Leica SP8 X confocal microscope with a 63×, 1.4-NA oil-immersion objective (Harvard NeuroDiscovery Center). We imaged areas of M2 that contained all cortical layers, with optical sectioning of 0.5 μm. For analysis, we utilized machine learning software^123^ to segment DAPI-stained nuclei and to count the fluorescent puncta of hybridized probes. For large area image presentation (**Fig. 1b**, **Fig. 2g**, **Supp. Fig. 1c**), nuclei masks created in the analysis pipeline are displayed and pseudocolored according to the number of puncta contained within the mask.

### Indirect calorimetry

14-week-old C57BL/6J male mice were housed at 23 °C in a Promethion indirect calorimetry system (Sable Systems International) within a temperature-controlled cabinet. The mice were injected with vehicle, T3, or sobetirome as described above. Data collected include VO2, VCO2, physical activity beam breaks, food intake, and body mass. For the duration of the experiment, the mice had ad libitum access to Labdiet 5008 (3.56 kcal/g) and were maintained on a 12hr/12hr photoperiod with lights on from 0600/1800.

### Light-dark behavioral paradigm

The Light-Dark box arena (27.3cm x 27.3cm x 20.3cm, Med Associates, ENV-510S-A) was divided into two compartments: an uncovered area under direct light and a covered area separated with an opaque plexiglass structure (Med Associates, ENV-511). An opening (6cm x 5 cm) in the plexiglass separator allowed mice to move between the two compartments. Mice spent 10 minutes in the arena. Time spent in the light and dark areas was analyzed using the Activity Monitor software (Med Associates, version 7). Light-dark boxes were cleaned with ethanol before each mouse began its trial and allowed to dry. Animals were 11-12 weeks old at the time of the experiment. In experiments that consisted of multiple rounds with multiple conditions, the data for each round was normalized to the control cohort of mice in the same round. Outlier animals greater than 3 median absolute deviations (MAD) from the median were removed from analysis.

### Immunohistochemistry

Animals were anesthetized by isoflurane inhalation and trans-cardially perfused with cold 0.9% saline followed by 4% paraformaldehyde (PFA) in a phosphate buffered solution (PB) (0.081 mM Na_2_HPO_4_, 0.017 mM NaH_2_PO_4_, pH 7.2-7.4). Brains were harvested and post-fixed in 4% PFA overnight at 4C and preserved in 0.5xPB + 0.02% sodium azide. Brains were sliced coronally at 50 μm and slices were permeabilized with a blocking buffer containing 0.1% Triton-X, 6% normal goat serum (NGS, Abcam ab7481) in PBS for 2 hours at RT. Primary antibodies were applied at 4° C overnight. Secondary antibodies were applied for 2 hours at room temperature and counterstained with DAPI (Sigma Aldrich D9542, 1:10000) for 20 minutes. Slices were then mounted to glass slides and coverslipped with Vectashield Vibrance (Vector H-1700) Primary antibodies used for immunohistochemistry (IHC) are listed here with dilutions indicated in parentheses: rat anti-HA (Roche 11867423001, 1:500), chicken anti-GFP (Abcam ab13970 1:1500), rabbit anti-NeuN (Sigma Aldrich ABN78). Fluorophore-conjugated secondary antibodies for IHC: goat anti-rat Alexa 555 (ThermoFisher A21434, 1:500), goat anti-chicken Alexa 488 (ThermoFisher A11039, 1:500), goat anti-mouse Alexa 647 (ThermoFisher A21235, 1:500), streptavidin Alexa 647 (ThermoFisher S32357, 1:50).

### Robo3 protein extraction and quantification

*Robo3^fl/fl^* mice were retro-orbitally injected with a systemic AAV serotype (PHP.eB) capable of efficiently transducing a Cre transgene to neurons in cortex (AAV_PHP.eB-Cre-GFP, 5 x 10^11^ vg). We then waited ∼3 weeks to allow for Cre-mediated excision of *Robo3* and subsequent protein turnover. Animals, C57BL/6J, un-transduced *Robo3^fl/fl^*, and Cre-transduced *Robo3^fl/fl^*, were treated with T3 or vehicle control. After 3.5 days of treatment anterior cortex was dissected and submerged in ice-cold 300 µl of RIPA buffer (Thermo Fisher Scientific) supplemented with protease inhibitors (cOmplete mini, Roche). The tissue was lysed in a Dounce homogenizer with 40 strokes of a plunger, and the resultant lysate was incubated with end-over-end rotation for 10 minutes at 4 °C. The lysate was then cleared by centrifugation (17000g, 10 minutes, 4 °C), and the supernatant was passed through a Wizard column to remove genomic DNA (Promega) (10000g, 1 minute, 4 °C). Protein from lysates were denatured by the addition of 50 ml of Laemmli sample buffer (37 mM Tris-HCl, pH 6.8, 10% (wt/vol) SDS, 25% (vol/vol) 2-mercaptoethanol, 25% (vol/vol) glycerol, and 0.056% (wt/vol) bromophenol blue). Samples were resolved by 8–16% SDS-PAGE, transferred for 2 hours at room temperature at 45 V to 0.45 mm PVDF membranes, and analyzed by immunoblotting. Membranes were blocked with 5% milk prepared in TBST (Tris-buffered Saline with Tween 20) for at least 5 min at room temperature, then incubated with primary antibodies in 5% BSA TBST overnight at 4 °C with end-over-end rotation. Primary antibodies targeting the following proteins were used at the indicated dilutions and obtained from the denoted companies: Robo3 1:300 (R&D systems, #AF3155), GluN1 NMDAR 1:1000 (SySy #114011). Following overnight incubation, membranes were washed three times, 5 min each, with TBST and incubated with the corresponding secondary antibodies in 5% milk (1:5000) for 1 hour at room temperature. Membranes were then washed three more times, 5 min each, with TBST before being visualized using enhanced chemiluminescence (Thermo Fisher Scientific). Signals were quantified using ImageJ (Fiji).

### Generation of single-nucleus suspensions from mouse secondary motor cortex

Fresh-frozen mouse brains were securely mounted by the cerebellum onto cryostat chucks using OCT embedding compound without thermal perturbation. Bilateral dissection of the anterior regions of secondary motor cortex (M2) was performed by hand in the cryostat using an ophthalmic microscalpel (FEATHER Incision scalpel P-715) and 4x surgical loupes. Each dissected sample was placed into a 0.25 mL PCR tube using forceps. All materials were pre-cooled to -20° prior to use.

Nuclei isolation was then performed as previously described^124,125^. Briefly, sectioned tissues were moved from the cryostat into 12-well plates (one well per sample), and 2 mL of extraction buffer was added to each well. Mechanical dissociation was performed by slowly triturating up and down 20 times using a P1000 pipette (1 mL Rainin tip), with extra care to avoid froth or bubbles. This trituration step was repeated three or four additional times, with a 2 minute break between rounds. Using a syringe, we passed each sample twice through a 26-gauge needle into its original well. ∼2 mL of this solution was transferred into a 50 mL conical tube for each sample, followed by the addition of wash buffer to a total volume of ∼20 mL. This mixture was then split equally into two 50 mL conical tubes. The samples were spun down in a swinging-bucket centrifuge at 600 RCF for 10 min at 4°C. After centrifugation, the supernatant was carefully removed, making sure not to disturb the pellet, until ∼500 µL remained in the tube. The resuspended nuclei were then pooled back into a single tube, resulting in ∼1 mL of concentrated nuclei solution. DAPI (Thermo Fisher Scientific, no. 62248) was added at 1:1000 concentration and incubated for at least 5 minutes prior to sorting. Singlet nuclei were isolated utilizing fluorescence activated cell sorting on a Sony SH800, and the final nuclei concentration was determined using a hemocytometer.

### snRNA-seq library preparation and sequencing

The 10X Genomics (v.3) kit was used for library preparation according to the manufacturer’s recommendation. Libraries were pooled and sequenced on either a NovaSeq S2 or NovaSeq S4. Sequencing reads were demultiplexed using the CellRanger v.5 pipeline and aligned to the GRCm38 reference genome, which was custom annotated to facilitate readout of Cre and DN/WT-THR expression.

### Major cell class annotation

Cell types were defined using a two-step process. First, we used a modified version of the Seurat v.2 workflow^126^ to perform initial quality control, clustering, and annotation of major cell classes. Briefly, nuclei with fewer than 200 total unique molecular identifiers (nUMIs) and greater than 5% mitochondrial genes (MT%) were excluded from subsequent rounds of analysis. The data were annotated to reflect the “genotype” (non-transduced C57BL6/J mice referred to as endogenous, virally transduced C57BL6/J with Cre and Cre-dependent DN-THR, or virally transduced C57BL6/J with Cre and Cre-dependent WT-THR) and treatment condition (T3 or vehicle control). Additional quality control metrics were calculated–including number of unique genes (nGene), percent oxidative phosphorylation genes (OXPHOS%), and percent ribosomal protein–and gene identifiers were mapped to Ensembl gene IDs. The virally transduced Cre and DN- or WT-THR transcripts were annotated prior to removal of any other genes lacking Ensembl gene IDs or with duplicate Ensembl gene IDs. Highly variable gene selection was performed using Seurat’s vst method to identify the top 2000 genes with expression patterns that drive biological differences between cell types while minimizing batch effects.

After scaling the data, we performed preliminary principal component analysis (PCA; k=25) and Uniform Manifold Approximation and Projection (UMAP) dimensionality reduction. Louvain clustering was used throughout all analyses with clustering resolutions of 0.6 or 1.0. Clusters with high MT% and OXPHOS%, which also had no clearly delineated cell type markers, were identified and removed from downstream analyses. A second round of PCA (k=100) and UMAP was performed on the filtered dataset and the resulting clusters were annotated into one of seven major cell classes (glutamatergic neurons, GABAergic neurons, astrocytes, oligodendrocytes, oligodendrocyte precursor cells, immune cells, and endothelial cells) based on per-cluster expression of a list of marker genes^47^. Putative doublets (clusters that showed substantial expression of marker genes from two or more major cell types) were removed.

### Subtype annotation based on mouse motor cortex reference dataset

Subtypes within each broad cell class were assigned using a random walk algorithm^126^ to transfer annotations from a recently published transcriptomic atlas of the mouse motor cortex^47^ (the “source” dataset) to our study dataset (the “target” dataset). For glutamatergic and GABAergic neurons, the original embedding space was sufficient for obtaining a high-quality annotation of clustered subtypes due to limited sample-to-sample batch effects and high similarity to the source dataset. For glial cell types (except oligodendrocyte precursor cells for which there was only one cluster), each cell class specific subset of our target dataset was first aligned to the corresponding cell types of the source dataset by performing an integrative analysis including highly variable gene selection and PCA (k=30 for oligodendrocytes and endothelial cell classes; k=100 for astrocytes and immune cell classes). We utilized Harmony to account for batch effects between the source and target datasets and to flag additional doublet clusters revealed by this cell class specific process for removal. Several immune cell clusters from the DN- and WT-THR genotypes failed to align with the source dataset–likely reactive microglia responding to viral infection– and were excluded from subsequent DE analyses. Subtype annotation was verified with per-cluster marker analysis.

### Pseudocell-based differential expression analysis

We utilized a previously developed pseudocell strategy^126^ coupled with linear modeling to identify differentially expressed (DE) genes. Within each cell subtype, we constructed pseudocells by aggregating the raw UMI count of, on average, 30 nuclei per sample. Each resulting pseudocell was thus composed of nuclei from mice of the same genotype (endogenous, DN-THR, or WT-THR) and treatment status (T3 or vehicle control). Pseudocells had a minimum of 15 nuclei, and cell subtypes with fewer than 15 nuclei or 6 total constructed pseudocells were excluded from analysis.

For all comparisons, we implemented the Limma Trend approach with robust moderated t-statistic and Benjamini-Hochberg (BH) corrected p-values to identify DE genes for each cell subtype. %MT and log_2_(nGene), calculated at the pseudocell level, were used as covariates and sample ID was used as a random effect. Genes expressed in greater than one percent of cells of the relevant subtype were used as background for the differential expression testing.

We first used the above DE methodology to assess the effect of thyroid hormone on gene expression in the motor cortex (**Fig. 2**, **Supp. Fig. 2-3**). Across a subset of pseudocells from “endogenous” samples, a DE analysis was performed between T3- and vehicle control-treated samples. A robustness score (rob.score; defined for each gene as the fraction of sample pairs in the experiment showing up-versus down-regulation based on the mean expression across the tested condition) and robustness percentage (rob.pct; defined similarly but using percent non-zero expression) was calculated for each gene. A TRG was then defined as any DE gene with BH corrected p-value < 0.05 and a robustness score of ≥ 0.5, meaning that a gene is consistently up- or down-regulated in at least 75% of comparisons. Non-TRGs (**Fig. 3i**, **Supp. Fig. 5**) were defined by subsetting the top 10,000 most expressed genes with a BH corrected p-value > 0.05, a rob.score of 0, and a rob.pct of 0.

A second set of DE analyses were performed (**Fig. 3**, **Supp. Fig. 5**) to evaluate the effect of DN-THR or WT-THR expression on the TRG programs identified above. New pseudocells were constructed by sorting nuclei according to their genotype (DN-THR or WT-THR only), treatment, and Cre status. Therefore, each pseudocell was composed of exclusively “Cre positive” nuclei– defined as having 1 or more detected Cre transcript(s) as a readout of the presence of activated Cre-dependent DN-THR/WT-THR constructs–or “Cre negative” nuclei–defined as having 0 detected Cre transcripts. DE analyses were performed to compare the Cre+ and Cre-conditions.

Given sample size limitations for the DN-THR/WT-THR experiment (2-3 mice per genotype), robustness scores were not calculated.

### TRG correlation matrix

To produce the correlation matrix in **Fig. 2D**, the Spearman correlations between log_2_(fold-change) values were calculated across all TRGs for each pair of cell types. Hierarchical clustering was then performed on the Euclidean distances of this TRG correlation matrix by implementing Ward’s clustering criterion (“ward.D2”). Only cell types that induced TRGs were included in this analysis.

### Ordered gene set enrichment analysis

For each cell subtype analyzed through the above differential expression framework, the background genes used in the DE analysis (genes expressed in >1% of cells of the relevant subtype) were annotated using the gene biotype information in the “EnsDb.Mmusculus.v79” package in R Bioconductor and filtered to include only protein-coding genes. These resulting genes were then ordered by the absolute value of the t-statistic derived from the Limma Trend model, such that the most significantly perturbed genes–regardless of the direction of the perturbation–were highest in the ranked list. The 2023 GO Biological Process gene set was obtained from the EnrichR portal^127^ and the fGSEA package v1.24.0^128^ was run with a “pos” scoreType, minimal gene set size of 15, and maximal gene set size of 500. To identify top fGSEA terms for each subtype (Fig. 2E), we subsetted significantly enriched pathways (BH-adjusted p-value<0.05) in which the leading-edge gene subset (i.e. the genes driving the enrichment signal) contained at least 3 TRGs, as defined in the endogenous T3 vs control DE analysis described above. The union of the top 3 fGSEA terms, ranked by BH-adjusted p-value, were included for each neuronal cell subtype.

### Stereotaxic surgeries

Mice were anesthetized with 2-3% isoflurane and 0.08% oxygen, and surgeries were performed under aseptic conditions within a stereotaxic frame (David Kopf Instruments). For intracranial injections, small craniotomies were drilled using a #81 drill bit (Kyocera, 20-517-025) and injections were performed through a pulled glass pipette containing virus, driven by a syringe pump (Harvard Apparatus, #883015) at a rate of 40 nL/min. After injections, the wound was sutured, mice were placed in a cage with a heating pad until their activity recovered, and then they were returned to their homecage and were given pre- and post-operative oral carprofen (CPG, 5 mg/kg/day) and monitored daily for at least 4 days post-surgery. Experiments were performed at least 13 days after viral injections to allow for transgene expression.

For 2ABT experimental animals, the same surgical setup was used to install a headpost. Briefly, the skull was lightly scored with a scalpel and a metal headpost was glued at lambda (Loctite gel #454). White cement (Flow-It ALC) was used to secure a border between the skin and the skull. For animals included in 2ABT experiments described in **Fig. 6**, a drill bit was used to lightly mark coordinates for future viral injections. The remaining exposed skull was covered with silicon (KwickKast). Animals were given CPG (5 mg/kg/day) and monitored as above before beginning 2ABT training.

The following coordinates were used as injection sites (relative to bregma):

**Fig. 3**, **Supp. Fig. 5**: +2.4 mm and +1.8 mm A/P, +/-1.0 mm M/L, and 0.6 mm D/V.

**Fig. 4, Supp. Fig. 7-8**: +2.2 mm A/P, +/-1.0 mm M/L, and 0.4 mm D/V.

**Fig. 6, Supp. Fig. 11-13**: +2.5mm A/P, +/-1.5mm and +/-0.5mm M/L, 0.4 and 0.9 mm D/V.

The following viruses were used for injections (final titer in gc/ml):

**Fig. 3**, **Supp. Fig. 5**: 400 nL scAAV2/9-hSyn-iCre-HA (9 x 10^10^) combined with either AAV2/9-SIO-nEF-DN-Thrb (5 x 10^12^) or AAV2/9-SIO-nEF-WT-Thrb (5 x 10^12^).

**Fig. 4**: Left hemisphere: 200 nL AAV2/9-hSyn-FLEX-CoChR-GFP (5 x 10^12^). Right hemisphere: 200 nL AAV2/retro-CAG-Cre (5 x 10^12^).

**Supp. Fig. 7**: Left hemisphere: 200 nL AAV2/9-hSyn-FLEX-CoChR-GFP (3 x 10^12^) combined with either AAV2/9-SIO-nEF-DN-Thrb (3 x 10^12^) or AAV2/9-SIO-nEF-WT-Thrb (3 x 10^12^). Right hemisphere: 200 nL AAV2/retro-CAG-Cre (5 x 10^12^).

**Supp. Fig. 8**: Left hemisphere: 200 nL AAV2/9-hSyn-CoChR-GFP (1 x 10^12^) combined with either scAAV2/9-hSyn-iCre-HA (1 x 10^12^) or scAAV2/9-hSyn-ΔiCre-HA (1 x 10^12^). Right hemisphere: no virus.

**Fig. 6, Supp. Fig. 11-13**: 200 nL AAV2/9-hSyn-DN-THR (1×10^13^), or 200 nL AAV-hSyn-WT-THR (1×10^13^).

### Electrophysiology

Brain slices were obtained from 2.5- to 4.5-month-old mice using standard techniques. Mice were anaesthetized by isoflurane inhalation and perfused trans-cardially with ice-cold ACSF containing (in mM) 125 NaCl, 2.5 KCl, 25 NaHCO_3_, 2 CaCl_2_, 1 MgCl_2_, 1.25 NaH_2_PO_4_, and 11 glucose (295 mOsm/kg). Brains were blocked, cut along the midline, and hemispheres with contralateral projecting CoChR+ axons were transferred into a slicing chamber containing ice-cold ACSF. Coronal slices of anterior cortex were cut at 300 μm thickness with a Leica VT1000 S vibratome in ice-cold ACSF, transferred for 10 min to a holding chamber containing choline-based solution (consisting of (in mM): 110 choline chloride, 25 NaHCO_3_, 2.5 KCl, 7 MgCl_2_, 0.5 CaCl_2_, 1.25 NaH_2_PO_4_, 25 glucose, 11.6 ascorbic acid, and 3.1 pyruvic acid) at 34°C then transferred to a secondary holding chamber containing ACSF and maintained at room temperature (20–22°C) until use. All recordings were obtained within 6 hours of slicing. Both choline solution and ACSF were bubbled with 95% O2/5% CO2.

Acute coronal slices were maintained in ACSF at 34 °C. Whole cell recordings were obtained from L2/3 glutamatergic neurons (150-350 μm below the pia surface) at ∼2.5-2 mm anterior to bregma, and ∼1 mm lateral of the midline. identified by morphological (pyramidal cell body) and electrophysiological (membrane capacitance C_m_ > 75 pF, membrane resistance R_m_ < 300 MΩ) features. For recordings of optically evoked PSCs, pipettes were pulled to have a resistance of 2-3 MΩ and were filled with internal solution containing (in mM), 135 CsMeSO_3_, 10 HEPES, 1 EGTA, 3.3 QX314-Cl, 4 Mg-ATP, 0.3 Na-GTP, 8 Na_2_-phosphocreatine. In a subset of experiments, 1 mg/ml biocytin (Sigma B4261) was added for post-hoc staining. Whole cell recordings were conducted in voltage-clamp mode, first at -70 mV to record evoked EPSCs, then at 0 mV to record evoked IPSCs. Full field illumination (0.13 mm^2^) was delivered by a 473 nm laser whose power output was controlled through an acousto-optic modulator. A 2 ms light pulse was delivered at randomized powers (7-10 discrete powers delivered between approximately 0.03 to 4 mW), with 15 seconds between pulses to allow for recovery. Recordings of each light stimulus were 3 s long, with a 1 s baseline prior to light pulse delivery. Each light power was repeated 3 times.

Quality control criteria were 1) manual inspection of unlabeled recordings for baseline stability (blinded to condition), 2) generation of peak currents > 250 pA at saturating light stimuli (> 3 mW) to indicate that recordings were made within the CoChR-labeled axonal field, 3) variation in Rm < 20% between recordings 4) holding current I_h_ < -400 pA at V = -70 mV, and 5) variation in series resistance Rs < 25% between recordings and V = -70 mV and V = 0 mV recording epochs. All analyses and figures use the 10 ms integrated charge at each power, normalized by C_m_. These values were fit in a custom MATLAB script to a sigmoid function, 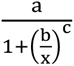, where *x* is the laser power, to find optimal values for coefficients. For figures, the log_2_(fold-change) value of an individual coefficient relative to the median control condition coefficient value is presented.

For measurements of intrinsic excitability, L2/3 glutamatergic neurons were targeted as above. Pipettes were filled with a potassium-based internal solution containing (in mM), 135 KMeSO_3_, 3 KCl, 10 HEPES, 1 EGTA, 0.1 CaCl_2_, 4 Mg-ATP, 0.3 Na-GTP, 8 Na_2_-phosphocreatine. Recordings were made in current-clamp configuration. Resting membrane potential (RMP) was first recorded, and then adjusted to -80 mV. There was no difference in recorded RMP between T3 and C (-87 ± 4 mV, -85 ± 5 mV, p = 0.06, Mann-Whitney U-test). 1 s current injections were delivered in 50 pA increments between -100 to 650 pA in random order., with an inter-pulse interval of 10 s. Recordings of each current injection were 2.5 s long, with a 1 s baseline prior to current injection. Experiments were repeated using alternate divalent cation concentrations in the ACSF, identical to the above except 1 mM CaCl_2_ and 2 mM MgCl_2_. Quality control criteria were a stable baseline membrane resistance (variation in R_m_ < 10%) and a stable baseline resting potential with median standard deviation of ≤ 0.5 mV across recordings.

### 2ABT

Mice had a headpost installed as described above. After recovery, mice were water restricted to 1 mL per day for five days prior to training. Mice were maintained at >80% initial body weight for the full duration of the training. Mice were age 7-11 weeks at the start of water restriction.

The behavior apparatus was contained within a sound-attenuating box (Med Associates ENV-018V) and consisted of a custom-built two-tiered platform. Two steel spouts (0.05” OD, 0.033” ID), 0.5 cm apart, were mounted on a 2-axis motor stage (Zaber A-MCB2-KS10A) via a post equipped with a manual 3-axis fine positioning stage (Narishige U-3C). Animals were located on the top tier of the platform on a detachable stage. The stage consisted of a copper lined, clear, polycarbonate round tube (3.8 cm ID x 10 cm, McMaster-Carr) with a 2 cm long, 3/4 cylinder opening, fixed to a circular aluminum breadboard (15 cm x 0.127 cm, Thorlabs MBR6) via an adjustable-height optics clamp (Thorlabs VG100). Custom metal headpost holders were secured to the stage, directly flanking the small opening of the tube, using two hex-locking post holders (2.5 cm, Thorlabs PH1) and mounting bases (2.5 cm x 5.8 cm x 1 cm, Thorlabs BA1s). The stage was secured within the behavior apparatus using kinematic bases with magnetically coupled plates (7.6 cm x 7.6 cm, Newport BK-3A). Speakers (Audax TW025A20) were mounted on the lower tier of the platform to provide auditory cues. Solenoids (Lee LHQA0531220H) were used to deliver water droplets.

The behavior task was run using custom software (LabVIEW 2014, National Instruments) utilizing a MyRio-1900 (National Instruments). The trial structure consisted of an enforced no-lick period (1-2 s), were any lick resulted in a trial restart, and a tone (75 ms long, 5 khz) marking the beginning of a selection period (3 s). During this period, a mouse could make a choice by licking one of the two spouts. Probability of water delivery was assigned in software. If successful, 2.5 ul of water was delivered from the selected spout during a 3 s consumption period (which occurred regardless of outcome). Trials were delineated into blocks in which one spout has a higher probability of reward than the other. At the end of each block period, the reward probabilities reversed.

Mice were first trained on a deterministic version of the task to teach switching between spouts. For each trial, one of the two spouts delivered a water droplet. The block length in the deterministic task was 8-9 rewards. Spout probabilities reversed after mice received the allotted number of rewards. After 3-7 days, mice were introduced to a probabilistic task with a block length of 20 trials. One spout within a block had a 90% probability of delivering a water droplet while the alternate spout had a 10% probability (90/10). These probabilities reversed with each block. After 1-8 more days of training, the reward probabilities were changed to 80% for one spout and 20% for the alternate spout within a block (80/20). For the cohort of wildtype animals in **Fig. 5c-i**, this was the final task structure. For the cohort of WT-THR or DN-THR injected animals in **Fig. 6**, we altered the block structure such that the block length varied randomly from 20-40 trials with an approximate exponential distribution (43% probability of a block of 20 trials in length, 32% probability of a block of 30 trials in length, and 25% probability of a block of 40 trials in length).

Performance was evaluated by probability of selecting the highly rewarding spout (p(high)) averaged over 5 days of training. Performance criterion was met by achieving a 5-day window with average p(high) > 0.6. For the cohort of wildtype animals described in **Fig. 5c-i**, animals began experiments once meeting performance criterion and having trained at 80/20 for 10 or more days. Mice were treated with vehicle solution (described above) for 3-5 days to establish baseline behavior (habituation sessions), and then were treated with T3 or continued vehicle administration for 8 days (days 0-7 of treatment). For the cohort of WT-THR or DN-THR animals in **Fig 6**, C57BL6/J mice were given free access to water after reaching performance criterion at 80/20, and intracranial injections of AAVs encoding either WT-THR or DN-THR were performed (described above). After post-operative recovery, mice were water restricted again and reintroduced to the task. After at least 13 days post-surgery, to allow for WT- or DN-THR expression, mice began the experiment, with 4-6 habituation sessions followed by 8 days of T3 treatment (days 0-7 of treatment). After completion of the final day of the task, most mice were perfused as described above (immunohistochemistry) for brain-wide mapping of DN-THR or WT-THR. For analyses comparing before and after treatment, habituation sessions were compared to sessions from days 4-7 of treatment from individual mice. For mice with more than 4 habituation sessions, the last 4 habituation sessions were used to ensure the datasets were balanced.

### 2ABT behavioral modeling

To model mouse choice behavior in the 2ABT we used Q-learning, a reinforcement learning framework that estimates values of each potential action decision and uses these values to generate predicted choices. In this case there are only 2 action values, one for selecting the left spout, Q_l_, and the other for selecting the right spout, Q_r_. These values evolve according to (see **Supp. Fig 10**):

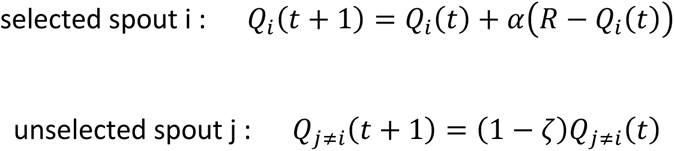

where R is the state of the water reward (R = 1 for a dispensed water droplet, R = 0 for a failure), α is the learning rate that scales the value update of the selected spout, and ζ is the forgetting rate which decrements the value of the unselected spout. Q-values are then fed into a softmax function that determines the probability of selecting each spout:

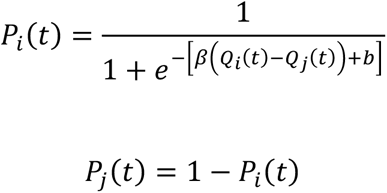

where β is an inverse temperature parameter given relative values between the spouts Q_i_(t)-Q_j_(t) and b is a bias parameter. β scales the slope of the sigmoidal activation curve and varies the degree to which actions are stochastic and exploratory versus exploitative of current value estimates. Models were fit to individual mice for habituation sessions, and separately for sessions from days 4-7 of treatment. Each session was split into 4, and model training was performed on randomly selected 3 of the 4 blocks of trials (75%), with the remaining block used as held out data to evaluate fits (25%). Parameters β, α, ζ, b were estimated using stochastic gradient descent optimization (learning rate of 0.1, 10000 iterations) and a negative log-likelihood loss function (custom Python code using PyTorch library). All fits were evaluated on the 25% held out data (**Fig. 5i, Supp. Fig. 13d**).

### Brain-wide mapping of DN-THR and WT-THR expression

Both DN-THR and WT-THR constructs have a C-terminal HA-tag to enable unambiguous quantification via using immunohistochemistry and imaging. Whole brain 50-micron slices were stained for the HA-tag and DAPI (described above) and were mounted and imaged using an Olympus VS200 Slide Scanner. NeuroInfo (Version 2020.1.1, 64 bit) was used to load and realign each image using the built-in BrainMaker Workflow. The Detect Cells feature was used to quantify cell positions in the anti-HA fluorescence channel (TRITC). Cells were identified based on size and intensity cutoffs calibrated to each brain. Cutoffs were determined by comparing manual annotations in a small area with NeuroInfo cell detection. Cell positions were then mapped to brain regions using the Allen Institute 2017 adult mouse common coordinate framework (CCF) atlas (25 μ/pixel) and counts per brain region were exported to a csv file. Mapped cell positions were exported from NeuroInfo into MATLAB, and a 3D Gaussian smoothing kernel (sigma = 5 pixels) was applied before cross sections were isolated and overlayed onto the Allen Mouse Brain CCF Atlas to visualize WT- or DN-THR expression as a heatmap using the imoverlay package (Matt Smith, version 1.3.0.0).

### Statistical analyses

Statistical tests used are listed either in the main text or figure captions. A description of the statistical methods used for differential gene expression are described above in the snRNAseq section. Likelihood ratio tests were performed on full versus reduced mixed effects models (MM) as follows: in **Fig. 1c** full models for energy expenditure, food consumption, and locomotion used animal weight, light/dark period, treatment, experimental time, and the interaction of treatment and experimental time as fixed effects, and individual animals as random effects. Reduced models included all the same terms, but without an interaction term. For energy expenditure we used a linear MM (LMM). Food consumption and locomotion were sparse datasets, so we used generalized linear MMs using zero-inflated negative binomial distributions. In **Supp. Fig. 1e** the full model for temperature (LMM) used animal age, treatment, experimental time, and the interaction between treatment and experimental time as fixed effects and individual animals as random effects. The reduced model included all the same terms but without an interaction term. In **Supp. Fig. 6**, full models (LMM) used the injected current and treatment as fixed effects, and individual cells as random effects. Reduced models lacked the treatment term. In **Fig. 5c**, the full model (LMM) used experimental day, treatment, and the interaction of treatment and experimental day as fixed effects, and individual animals as random effects. The reduced model included all the same terms but without an interaction term. The full model in **Fig. 6c** used experimental day, “genotype” (intracranial injections DN-THR or WT-THR), and the interaction of “genotype” and experimental day as fixed effects, and individual animals as random effects. The reduced model included all the same terms but without an interaction term.

### Code and data availability

Data were analyzed using custom scripts in MATLAB (2021b), R (4.2.2), and Python (3.9). Code and data are available upon reasonable request. Upon manuscript acceptance, snRNAseq sequencing data will be uploaded to GEO and data objects will be uploaded to a public repository.

